# Thymic keratin 76 shapes skin-specific central tolerance

**DOI:** 10.64898/2026.08.27.747460

**Authors:** Mara Gelmetti, Inês M. Tomás, Roberta Ragazzini, Sara Campinoti, Megan S. F. Soon, Mafalda Cautela, Matteo Vietri Rudan, Marina Torre, Nital Sumaria, Inês Saldanha, Diana Pereira, Norman Yap, Mirjana Efremova, Zewen K. Tuong, Fiona M. Watt, Paola Bonfanti, Daniel J. Pennington, Inês Sequeira

## Abstract

Keratin gene mutations are often associated with inflammatory skin disorders, in which the ensuing immunopathology is generally attributed to disrupted barrier integrity and consequent microbial invasion. Here, we challenge this paradigm by demonstrating a role for keratin 76 (Krt76) in thymic central tolerance to skin-targeting autoimmune responses. We show that transfer of Krt76^-/-^ thymic lobes under the kidney capsules of athymic recipients is sufficient to induce expansion of effector T cells in the secondary lymphoid organs, T cell skin infiltration, and autoantibody reactivity to both skin and oral mucosa tissue. Mechanistically, we demonstrate that loss of thymic Krt76 expression disrupts canonical differentiation of the thymic medulla and impacts the development of post-AIRE-expressing keratinocyte-like mimetic medullary epithelial cells (termed CorneoTECs). Notably, Krt76-expressing CorneoTECs differentially express a specific skin and oral mucosa-associated gene signature, including skin-specific tissue self-antigens (TSAs). Importantly, in the absence of Krt76 this skin and oral mucosa TSA signature is almost entirely lost, and T cell negative selection is affected. Collectively, these data highlight a heretofore unanticipated role for Krt76 in thymic central tolerance to skin and oral mucosa-targeting T cells and suggest that loss-of-keratin-associated skin disorders could also include autoimmune pathologies.

## INTRODUCTION

Skin inflammatory disorders involve a complex interplay between genetics, immunology, and epithelial barrier biology. This is evidenced by mutations in keratin (KRT) genes, which include those responsible for epidermolysis bullosa simplex, epidermolytic ichthyosis, epidermolytic hyperkeratosis, and pachyonychia congenita^1–8^ that drive skin fragility, chronic inflammation, and abnormal keratinocyte proliferation. Indeed, while best known for their structural roles in skin integrity, keratins are now recognised for their wider impact on immune regulation^7,9–11^. As an example, aberrant expression of KRT6, KRT16, and KRT17 is implicated in the immunopathology of inflammatory skin diseases such as psoriasis and atopic dermatitis^12^.

Keratins are not restricted to skin. Notably, keratin expression is a prominent feature of the thymus, a primary lymphoid organ that facilitates thymopoiesis and is necessary for establishing central tolerance that prevents self-reactive T cells from driving dangerous autoimmune pathologies in peripheral tissues. As in skin, thymic keratin expression delineates functionally distinct compartments. In mice, *Krt8* is predominantly expressed by cortical thymic epithelial cells (cTECs) and at the cortico-medullary junction^13^, while *Krt5* and *Krt14* are largely expressed by medullary thymic epithelial cells (mTECs). Moreover, a subset of mature mTECs (termed corneoTECs), adopt a keratinocyte-like phenotype that resembles terminally differentiated skin keratinocytes with expression of *Krt1* and *Krt10*, alongside other skin-specific markers^14–16^. These corneoTECs are one of several recently described “mimetic” mTEC subsets, some of which differentiate from autoimmune regulator (AIRE)-expressing precursors to adopt characteristics of cells from peripheral tissues^14,15,17–21^. Among these are mTECs that mimic: ciliated cells, expressing *Foxj1*; ionocytes, expressing *Foxi1* and *Foxi2*; microfold cells, expressing *Spib* and *Sox8*; muscle cells, expressing *Myog*, tuft cells, expressing *Pou2f3*; and neuroendocrine cells, expressing *Foxa*^14,15^. This cellular mimicry permits thymic expression of an antigenic “shadow” of peripheral tissues, to facilitate negative selection of tissue-specific self-reactive T cells and establish central tolerance^22–24^.

Here, we identify keratin 76 (Krt76), an important structural component of skin^25^, as a key factor shaping skin-directed central tolerance. *Krt76*-deficient mice display lymphadenopathy and splenomegaly, and progressive skin inflammation characterised by immune cell infiltration and autoantibody production^11,26^. Thymic transplantation of *Krt76*^-/-^ thymic lobes under the kidney capsule of *Foxn1^nu^* (nude) mice demonstrated that these phenotypes were thymus-driven. Using genetic models and single-cell transcriptomics, we reveal that *Krt76* deficiency disrupts standard differentiation of medullary areas and expression of a skin and oral mucosa-specific Krt76-dependent transcriptional signature. Collectively, these findings position Krt76 as an active player in thymic mimetic mTECs biology and establishing central tolerance that targets skin-specific autoimmune responses.

## RESULTS

### Thymic loss of *Krt76* causes skin-specific immunopathology

We previously reported an inflammatory phenotype in *Krt76*^-/-^ mice^11^. Here, we extended this characterisation to postnatal day-28 (P28) animals, an earlier, previously unexamined age-point, selected to set an appropriate baseline for the thymic transplantation experiments. By P28, CD4^+^ and CD8^+^ effector T cells were already expanded in spleen and lymph nodes (LNs) of *Krt76*^-/-^ animals (**Fig. 1a,b**, **Supplementary Fig. 1a,b**), with a concomitant increase in CD39^+^CD73^+^ regulatory T cells (Tregs) (**Supplementary Fig. 1c**), yet no differences in thymocyte populations were detected (**Supplementary Fig. 1d**). Notably, these expansions were accompanied by significant infiltration of CD45^+^ cells into the skin of *Krt76*-deficient mice, which included both CD4^+^ and CD8^+^ T cells (**Fig. 1c, Supplementary Fig. 1**), also seen in the tongue but not the muscle, suggesting a selective tissue targeting of inflammatory responses in *Krt76*^-/-^ mice (**Supplementary Fig. 1f, g**).

**Fig. 1.**
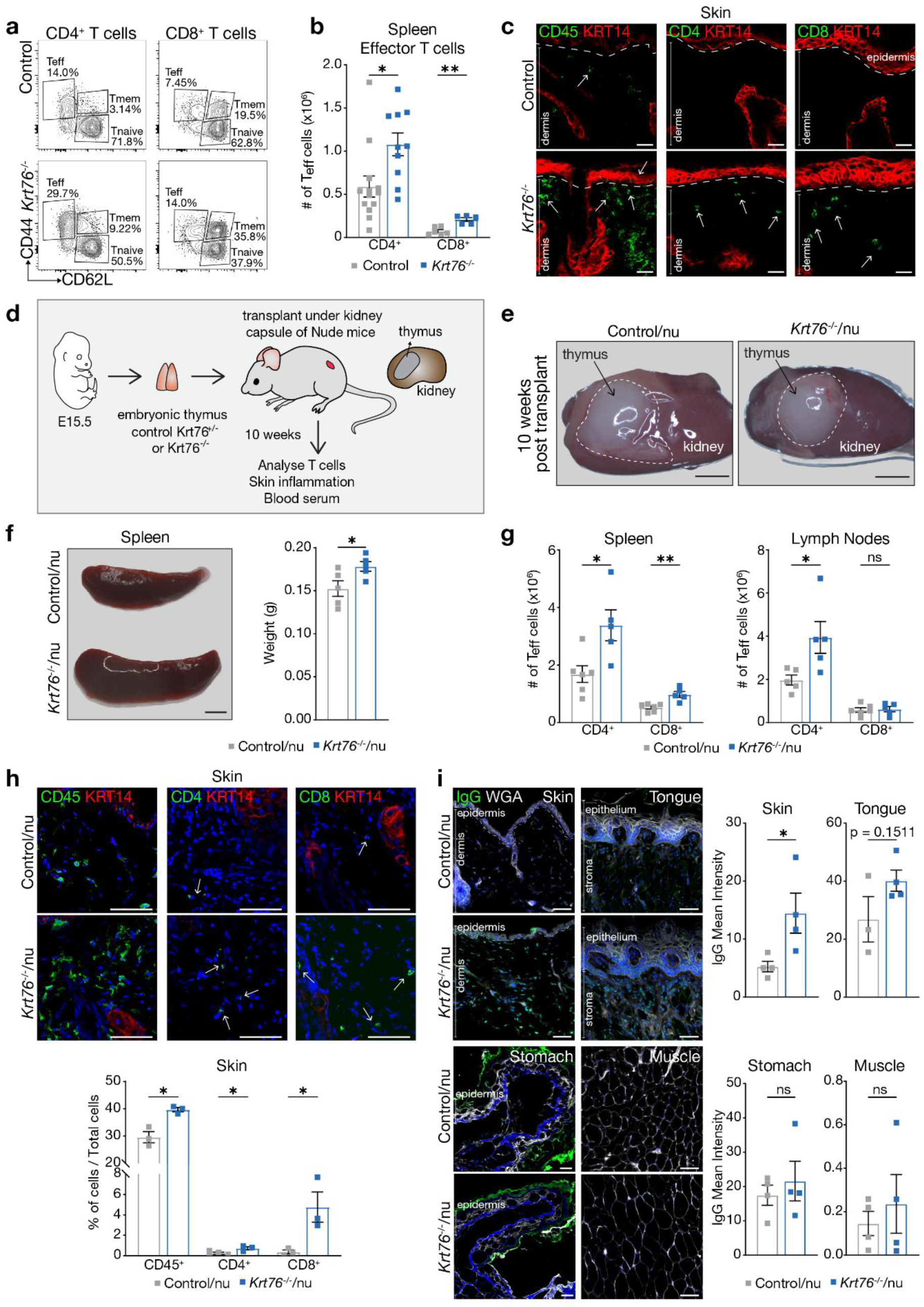
Loss of Thymic Krt76 instigates a T cell imbalance in the periphery. **a**, Representative dot-plots of CD44 and CD62L staining of postnatal day 28 (P28) splenic CD4^+^ and CD8^+^ T cells in Control and *Krt76*^-/-^ mice. **b**, Absolute number of splenic CD4^+^ and CD8^+^ effector T cells in Control and *Krt76*^-/-^ mice. **c**, Immunofluorescence staining of P28 Control and *Krt76*^-/-^ back skin for anti-CD45, anti-CD4 or anti-CD8 (green, arrows) and anti-KRT14, epithelial marker (red), **d**, Schematic workflow of renal capsular transplantation of thymi into Nude (nu) mice. E15.5 thymi from *Krt76*^-/+^ (Control) and *Krt76*^-/-^ grafted in the kidney subcapsule of adult nude mice. **e**, Representative photographs of Control/nu and *Krt76*^-/-^/nu thymi in Nude mice kidney subcapsule 10 weeks post-grafting. **f**, Representative photographs of spleen from grafted nude mice 10 weeks post-grafting, with spleen weight (g) quantification in Control/nu and *Krt76*^-/-^/nu mice. **g**, Flow cytometry analysis of spleen and lymph nodes absolute numbers CD4^+^ and CD8^+^ effector T cells for Control/nu and *Krt76*^-/-^/nu mice. **h**, Immunofluorescence staining of skin sections for either anti-CD45, -CD4 or -CD8 (green, arrows), anti-KRT14 (red) and DAPI (nuclear counterstain), with quantification of infiltrating CD45^+^/CD4^+^/CD8^+^ cells/mm^2^ in skin dermis versus DAPI^+^ total cells for Control/nu and *Krt76*^-/-^/nu mice. **i**, Immunofluorescence staining and quantification of IgG autoantibodies in sera of Control/nu and *Krt76*^-/-^/nu mice detected by staining *Rag1*^-/-^ mice back skin, dorsal tongue, squamous stomach and muscle sections, and measuring the mean fluorescent intensity. Wheat Germ Agglutinin (WGA) (white, membrane counterstain). Dotted lines delineate dermal-epidermal or stromal-epithelial junctions. Mean±s.e.m. t-test statistical analysis, \**p*≤0.05, \*\**p*≤0.01, \*\*\*\**p*≤0.0001. For IHC quantification, n≥3 sections/3-4 animals/group. Representative of *n*=2-5 experiments consisting of 3-13 animals/group. Scale bars: 10μm (c, h), 25μm (i), 1mm (f), 2mm (e).

Previous studies had suggested that skin inflammation in Krt76^-/-^ mice was a consequence of a skin barrier defect^25^. However, given that there was no epithelial barrier impairment in the tongue^11^, we considered that loss of *Krt76* may also affect immunological tolerance, leading to an auto-antigen driven response in the skin and oral mucosa. To provide evidence for this, we transplanted embryonic day-15 (E15.5) thymic lobes from *Krt76*^-/-^ or control animals under the kidney capsule of *Foxn1^nu^* (nude) athymic mice (**Fig. 1d**). Ten weeks later, grafted lobes from control/nu and *Krt76*^-/-^/nu mice looked comparable (**Fig. 1e**) and showed similar proportions and numbers of thymocyte populations (**Supplementary Fig. 2a**). By contrast, spleens from *Krt76*^-/-^/nu animals were notably larger and heavier (**Fig. 1f**), and there were increased numbers of CD4^+^ T cells in spleen and LNs, and CD8^+^ T cells in spleen (**Fig. 1g**), mirroring the phenotype of *Krt76*^-/-^ mice (**Fig. 1a,b**, **Supplementary Fig. 1a,b**)^11^. Back skin from Control/nu and *Krt76*^-/-^/nu mice did not show skin damage or changes to the epidermis thickness (**Supplementary Fig. 2b**). However, back skin of *Krt76*^-/-^/nu mice displayed an influx of CD45^+^ cells, which again included a significant increase of CD4^+^ and CD8^+^ T cells compared with skin from Control/nu animals (**Fig. 1h**). These data suggest that a thymus-dependent immunopathology develops in the absence of *Krt76*. Consistent with this, serum from *Krt76*^-/-^/nu mice, but not from Control/nu animals, contained autoantibodies that bound to skin and dorsal tongue sections, but not squamous stomach, muscle sections, oesophagus, heart, liver, kidney, pancreas and testis from Rag1^-/-^ mice (that do not produce antibodies) (**Fig. 1i, Supplementary Fig. 2c**). Thus, loss of Krt76 expression in the murine thymus results in an autoimmune pathology directed toward the skin and oral mucosa.

### Krt76 is expressed in distinct subsets of medullary thymic epithelial cells

To investigate further the role of KRT76 in establishing immunological tolerance in the thymus, we used single-cell RNA-sequencing (scRNA-seq), immunohistochemistry and flow cytometry to localise KRT76 expression in the thymus. *Krt76* expression appears at E18.5 and continues into adult age in mice (**Supplementary Fig. 3a, b)**, where it is expressed exclusively within the epithelial thymic compartment (**Supplementary Fig. 3c)**. Immunohistochemistry reveals KRT76 expression in mouse and human of the thymic medulla region that also expresses KRT14 (**Fig. 2a**). In human sections, KRT76 co-localised with involucrin and KRT10 within Hassall’s corpuscles that are spherical terminally differentiated structures formed of cells that resemble keratinocytes (**Fig. 2b**)^27^. These structures are not obviously present in the mouse thymus, yet KRT76 still co-localised with both involucrin and KRT10, markers of differentiated mTECs (**Fig. 2c**). While KRT76 is not expressed in all KRT10+ or Involucrin+ cells (**Supplementary Fig. 3d**), *Krt76* depletion reduces the proportion of double-positive cells (KRT10+KRT76+ or Involucrin+KRT76+) (**Supplementary Fig. 3e**). To assess this expression in greater detail, we utilised previously published human and mouse thymus scRNA-seq datasets^14,28^, in which we could identify and annotate several clusters of cortical thymic epithelial cells (cTECs) and mTECs. In both species, *Krt76* expression was predominantly restricted to the *Aire*^+^ mTECs II and post-*Aire* mTECs III clusters (**Fig. 2d**), confirmed via immunohistochemistry (**Supplementary Fig. 3f**). A recent seminal study demonstrated that mTECs can be further sub-clustered based on their mimetic characteristics, namely the expression of specific sets of tissue-restricted genes^14^. These include mTECs that mimic ciliated cells, ionocytes, microfold cells, muscle cells, entero/hepatic cells, tuft cells, endocrine cells, and corneocytes^14,15^. Thus, we re-analysed this mouse dataset^14^ to reveal that *Krt76* expression was largely confined to the differentiated keratinocyte-like CorneoTECs cluster that also expresses CorneoTECs markers *Krt1* and *Involucrin*^14–16^ (**Fig. 2e**). Consistent with this, KRT76 was not detected in polykeratin (PolyKRT) cells (**Fig. 2f**), a recently identified subset of epithelial stem cells in the human thymus^29^. However, upon induced differentiation of PolyKRT cells, KRT76 expression was upregulated (**Fig. 2f,g**), in a similar way to KRT7 and KRT10 which are associated with differentiated mTECs subtypes^29^. Collectively, these data demonstrate a highly restricted expression of Krt76 in both mouse and human thymus and suggest a role for KRT76 in mimetic mTECs that shape central tolerance to skin antigen-specific T cells.

**Fig. 2.**
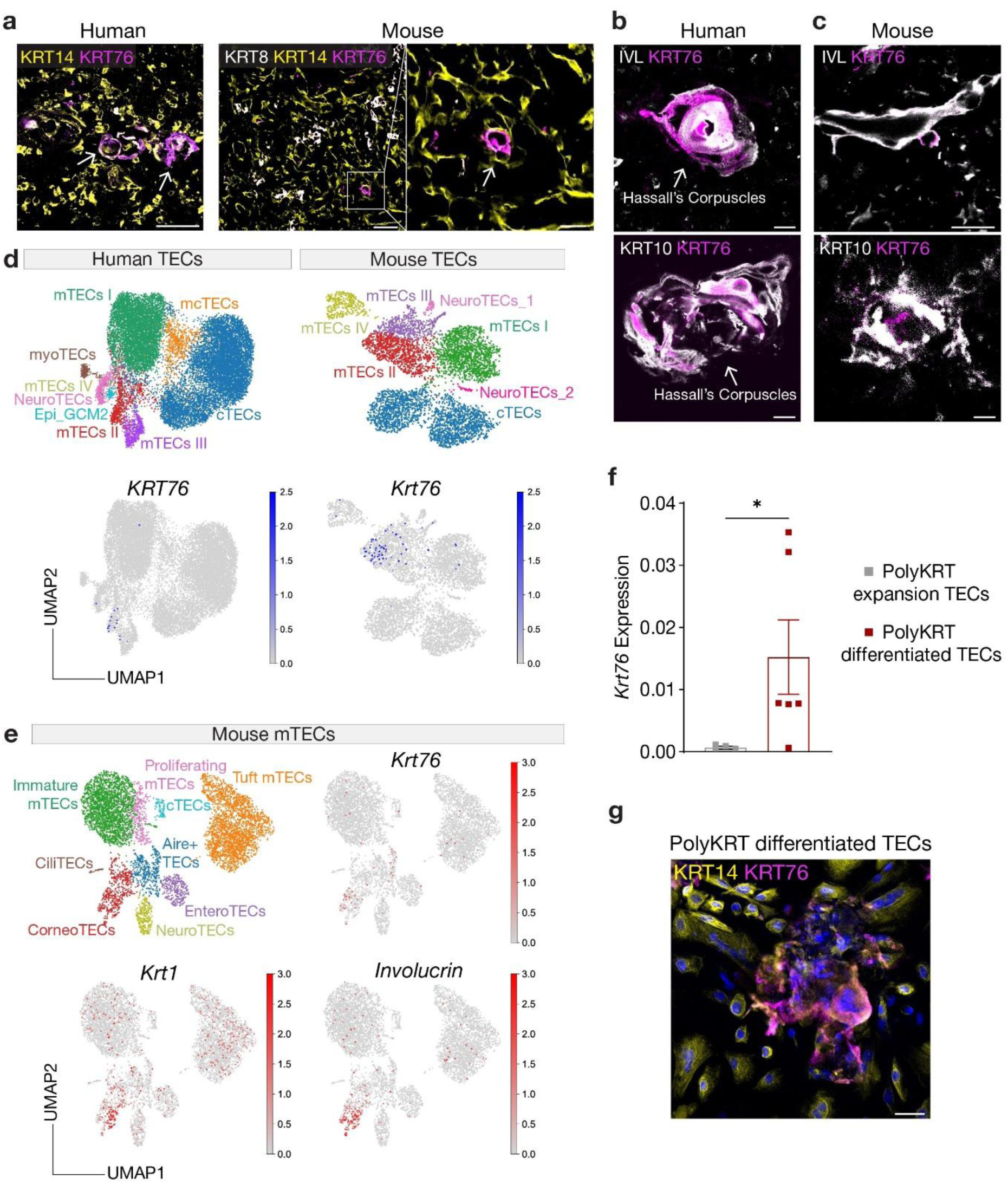
Terminally differentiated mTECs and Corneocyte-like Mimetic mTECs express KRT76. **a**, Immunofluorescence staining of human and adult C57/BL6 mouse thymus sections with anti-KRT8 (white), anti-KRT14 (yellow) and anti-KRT76 (magenta, arrows). **b**, Human and **c**, adult mouse thymi labelled with anti-KRT76 (magenta) and anti-Involucrin or anti-KRT10 (white). **d**, Uniform manifold approximation and projection (UMAP) of scRNA-seq thymic epithelial cell subpopulations in human and mouse datasets with *Krt76* expression (GEO: E-MTAB-8581^28^). **e**, UMAP visualisation of scRNA-seq Mimetic mTECs subpopulations of 4-6 weeks mice with *Krt76*, *Krt1* and *Involucrin* expression (GEO: GSE194252^14^). **f**, *Krt76* expression relative to HPRT in cultured human thymus PolyKRT (thymic BCAM^+^ stem cells)^29^ expanded TECs versus differentiated TECs. **g**, PolyKRT stem cells were cultured in expansion media (expanded PolyKRT TECs) for two days before differentiation assay gives rise to PolyKRT differentiated TECs. Immunofluorescence staining of PolyKRT differentiated TECs with anti-KRT14 (yellow), anti-KRT76 (magenta) and DAPI (nuclear counterstain). Mean ± s.e.m. t-test statistical analysis, \**p*≤0.05. *n*=6 animals/group. Scale bars: 10μm (b), 20μm (a), 50μm (a, f).

### Loss of Krt76 impacts mimetic mTECs differentiation

Having established that Krt76 is expressed in distinct thymic subsets of mimetic mTECs, the CorneoTECs, we next addressed the impact of loss of *Krt76* on mTECs differentiation. We sorted mTECs (CD45^-^EpCAM^+^Ly51^-^) from Control and Krt76^-/-^ mice and performed scRNA-sequencing to assess potential changes to mTECs biology in the absence of *Krt76*. A Tier-1 analysis generated 9 clusters that included 4,271 cells from Control thymus and 7,343 cells from Krt76^-/-^ thymus (**Supplementary Fig. 4a**). In addition to mTECs subsets, smaller clusters of cTECs, endothelial cells, fibroblasts and immune cells were present. We next performed a second level of clustering (Tier-2) on mTECs subsets only. This generated 12 clusters that we annotated as: immature mTECs I (expressing *Ccl21a*), proliferating mTECs (*Mki67*), *Aire*-expressing mTECs II (*Aire*, *Cd52*), and various subsets of recently described mimetic mTECs (mTECs III and mTECs IV). These included keratinocyte/CorneoTECs (*Spink5*), microfold mTECs (*Ccl20*), endocrine mTECs (*Cacna2d1*), entero/hepatic mTECs (*Lgals4*), ciliated mTECs (*Dynlrb2*), myocyte mTECs (*Myl1*), ionocyte mTECs (*Ascl3*), and tuft cell mTECs (*Trpm5*) (**Supplementary Fig. 4b**). We also identified *Krt5* and *Krt14* expression in mTECs I, mTECs II, and in proliferating mTECs, while *Krt1, Krt6a, Krt10, and Krt76* were all detected in mTECs II and corneoTEC clusters (**Supplementary Fig. 4b**).

Using this dataset, we assessed the impact of *Krt76* loss on mTECs differentiation. In comparison with Control mTECs subsets, *Krt76*^-/-^ mice displayed relatively higher proportions of early mTECs populations (mTECs I), and lower proportions of more differentiated mimetic mTECs subsets (**Fig. 3a**). Flow cytometry analysis of CD104 expression across mTECs subsets (CD104⁺ mTEC I vs. CD104⁻ mTEC III) validated our scRNA-seq observations (**Fig. 3b**). Additionally, thymus from *Krt76*^-/-^ mice show a decreased medullary representation of cells expressing KRT10, a marker of terminally differentiated keratinocytes (**Supplementary Fig. 5a**). Meanwhile, quantification of KRT14^+^- medullary regions across sequential thymic sections showed a reduction in proportional medullary areas, in medullary islets and in overall total medullary area (µm) in *Krt76*^-/-^ mice when compared to controls (**Supplementary Fig. 5b**), suggesting *Krt76* loss may contribute to impaired epithelial differentiation. Investigating more deeply, we used *scVelo*, which utilises the ratio of spliced and unspliced mRNA transcripts to predict the expected developmental trajectories of future cell transitions from our scRNA-seq dataset (**Fig. 3c**). In the *Krt76*-expressing clusters (*Aire*-high mTECs II, *Aire*-low mTECs II and CorneoTECs), Control mTECs displayed an organised set of trajectories that followed the established pathway from *Aire*-expressing mTECs stages to mimetic mTECs stages^14,30^. However, *Krt76*-deficient mTECs displayed a disorganised, disrupted set of trajectories, suggesting that loss of *Krt76* may disrupt CorneoTECs development, but not other Mimetic mTECs (EndoTECs, MicroTECs, Tuft mTECs) (**Fig. 3c**). Gene expression analysis across a continuous pseudotime trajectory showed an ordered expression of epithelial differentiation markers in control mTECs (**Fig. 3d**). By contrast, similar analysis of *Krt76*^-/-^ mTECs revealed disrupted expression of these genes, with delay of expression of genes important for the final stages of epithelial cell differentiation, such as Krt10 (*Krt10*), filaggrin (*Flg*) and Loricrin (*Lor*) (**Fig. 3d**). Thus, these data suggest that loss of thymic *Krt76* expression results in disrupted mTECs differentiation toward the mimetic mTECs stage, with particular disruption of the developmental trajectory that generates the skin-mimetic CorneoTECs population.

**Fig. 3.**
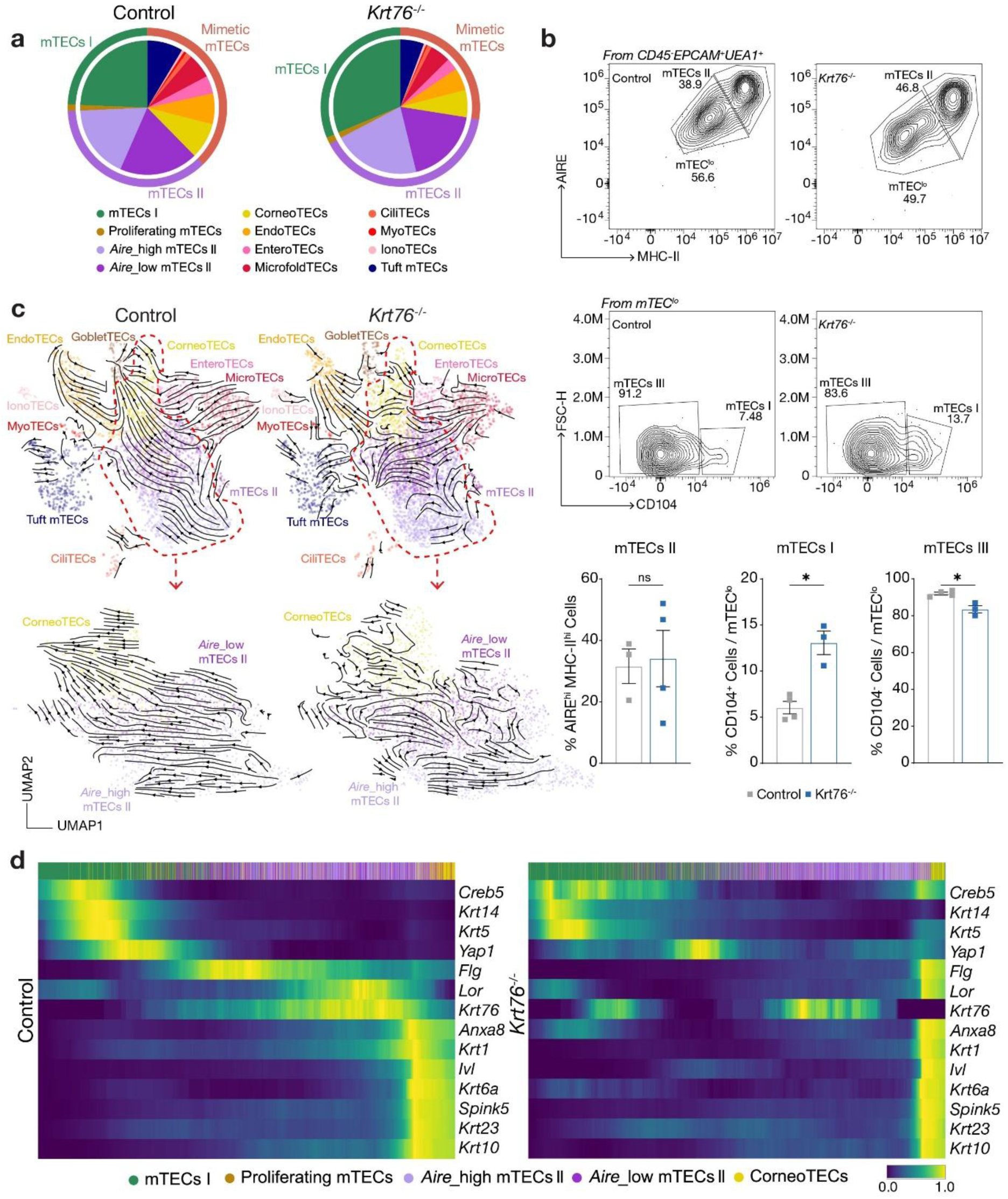
Loss of *Krt76* impacts mimetic cell differentiation. **a**, Pie chart of mTECs subsets proportions for Control and *Krt76*^-/-^ mice calculated based on scRNA-seq clustering annotation. **b,** Flow cytometry dotplots of mTEC^hi^ (AIRE^+^MHC-II^+^) (mTECs II) and mTEC^lo^ (AIRE^lo^MHC-II^lo^), and mTEC^lo^ CD104^+^ (mTECs I) and CD104^-^ (mTECs III) in 4-6 weeks Control and *Krt76*^-/-^ mice with representative quantification. **c**, Trajectory analysis of Control and *Krt76^-/-^* samples for mTECs II and Mimetic mTECs, along with *Krt76*-expressing clusters (*Aire*_low mTECs II, *Aire*_high mTECs II and CorneoTECs), dotted red line delineates *Krt76*-expressing clusters, arrows indicate forward direction. **d**, Heatmap representing diffusion pseudotime (DPT) expression of selected genes in Control and *Krt76^-/-^*samples across mTECs I, Proliferating mTECs, *Aire*_low mTECs II, *Aire*_high mTECs II and CorneoTECs subtypes. Mean±s.e.m. t-test statistical analysis, \**p*≤0.05. Representative of *n*=3 experiments consisting of 3-4 animals/group. *n*=2 scRNA-seq experimental repeats.

### Loss of *Krt76* downregulates expression of skin-related TSAs in mimetic mTECs

To further investigate the impact of *Krt76* loss on mimetic mTECs differentiation, we subclustered the scRNA-seq dataset, including only post-*Aire* mimetic mTECs (i.e. mTECs III and mTECs IV, Tier-3). This analysis yielded 12 clusters containing 1,770 control mTECs and 2,315 *Krt76*^-/-^ mTECs (**Fig. 4a**). Consistent with recent advances in mTECs characterisation^14,15,28–30^, we identified mimetic mTECs clusters resembling microfold cells, entero/hepatic cells, keratinocytes, goblet cells, endocrine cells, ciliated cells, ionocytes, myocytes, and tuft cells (**Fig. 4a, Supplementary Fig. 6a**). Differential gene expression analysis of control mTECs identified a *Krt76*-expressing mTECs-specific gene signature that included *Clic3*, *Gsta2*, *Defb6*, *Dapl1*, *Pinlyp*, *Trim29*, *Krt79*, *Aldh3b*2, *Dmkn*, *Sytl1*, *Il20rb*, *and Timp2* (for full list see **Supplementary Table 1**), many of which are associated with skin epithelia (**Fig. 4b)**. Notably, the expression of these genes is conserved in human TECs (**Supplementary Fig. 6b**), in both human and mouse skin scRNA-seq datasets^14,15,28–31^ (**Fig. 4b, Supplementary Fig. 7a,b**), as well as in human skin tissue sections^32^ (**Fig. 4c**). Examining the distribution of these *Krt76*-associated genes across our annotated control mTECs dataset revealed that many were enriched in CorneoTECs compared with the other mimetic mTECs subsets (**Fig. 4d**), although a subset (e.g. *Arhgef10l*, *Urah*, and *Timp2*) was also detected in other mimetic mTECs such as ionoTECs and myoTECs. Moreover, cross-referencing these genes with the Tabula Muris scRNA-seq atlas of all mouse organs^33^ showed that most of these genes had a restricted expression confined to skin and tongue (**Fig. 4e**). Thus, Krt76-expressing mTECs, that significantly overlap the mimetic CorneoTECs subset differentially express many genes associated with the skin and oral mucosa, the two tissues in which we observed immune cell infiltration and autoantibody binding in *Krt76*^-/-^ mice. Remarkably, when we assessed *Krt76*- expressing mTECs from *Krt76*^-/-^ mice (identified by detection of non-productive Krt76 transcripts), the expression of the TSA genes corresponding to the *Krt76*-associated gene signature that we had identified was significantly decreased and expressed by fewer cells (**Fig. 4f**). This finding reveals a major molecular consequence of *Krt76* loss.

**Fig. 4.**
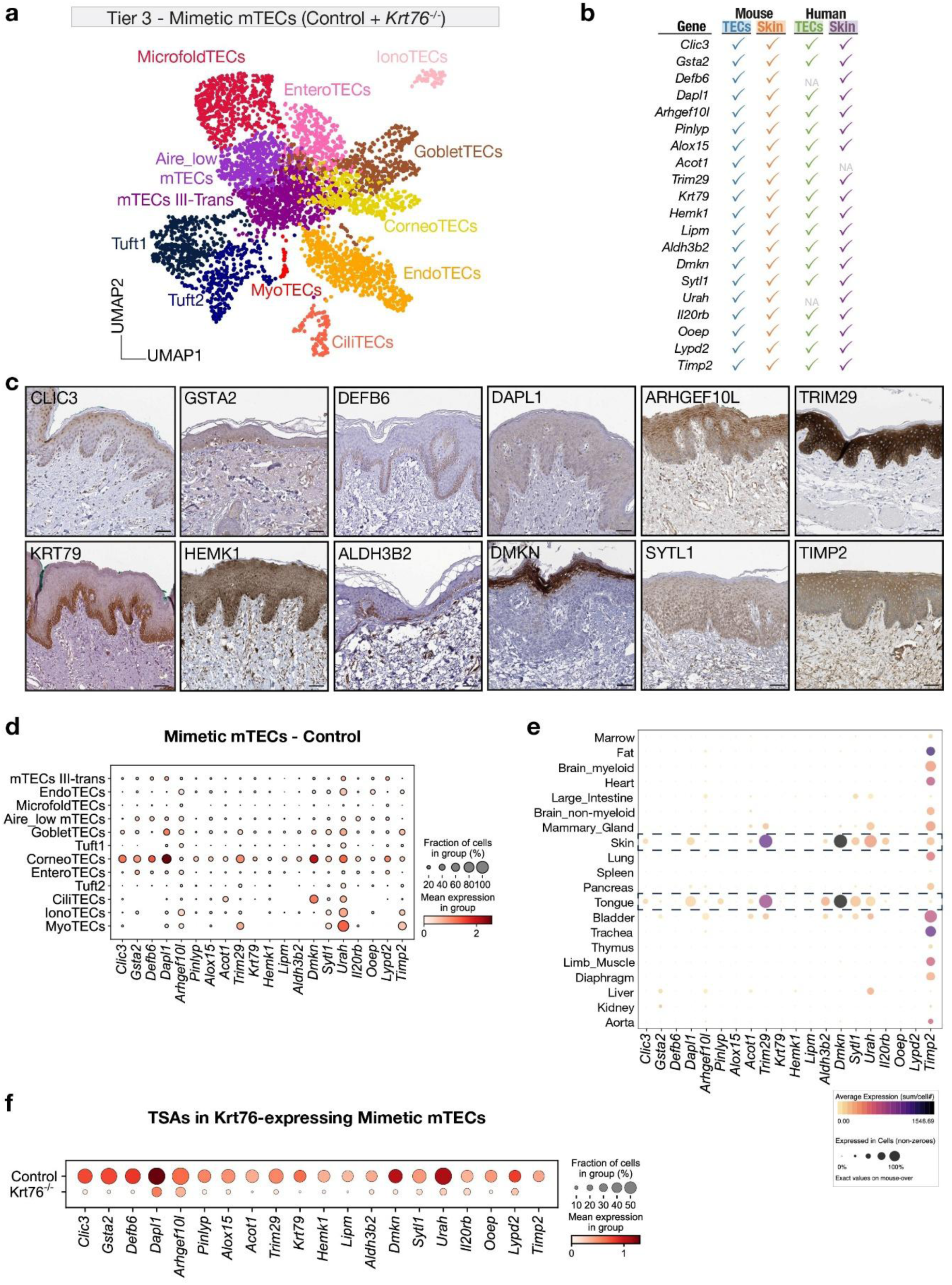
Loss of *Krt76* in Mimetic mTECs leads to downregulation of specific tissue self-antigens. **a**, scRNA-seq UMAP visualisation of Tier-3 subclustering of Mimetic mTECs subpopulations for integrated Control and *Krt76*^-/-^ mice (Control=1,770 cells; *Krt76*^-/-^=2,315 cells). **b**, Table summary of genes preferentially expressed in *Krt76*⁺ mTECs and their expression across Mouse and Human TECs and torso/dorsal Skin from public datasets (GEO: E-MTAB-8581, GSE129218, GSE241132)^28,31,51,52^, ✓ indicates gene expression present; NA indicates data not available. **c**, Immunostaining of CLIC3, GSTA2, DEFB6, DAPL1, TRIM29, KRT79, HEMK1, ALDH3B2, DMKN, SYTL1, TIMP2 expression in human skin sections, from the Human Protein Atlas^32^. **c**, Table of selected genes identified as *Krt76*- expressing TSAs along with a summary of their enriched tissue expression. **d**, Mean expression of genes identified as *Krt76*-expressing TSAs in all Mimetic mTECs clusters within the Control sample. **e**, Levels of expression of selected *Krt76*-expressing TSAs in all mouse organs. Data from the Tabula Muris Consortium, made available by the Chan Zuckerberg Biohub^33^. **f**, Mean expression of tissue self- antigens in *Krt76*-expressing Mimetic mTECs subpopulations for Control and *Krt76*^-/-^ mice (Control=36 cells; *Krt76*^-/-^=34 cells). Scale bars: 100µm (c). *n*=2 scRNA-seq experimental repeats.

To further characterise the candidates genes, we profiled serum from Control/nu and Krt76^-/-^/nu mice on the HuProt human proteome microarray. After excluding proteins with low mouse–human homology or lacking thymic-restricted expression (TRE) annotation^24^, CLIC3 emerged within the top 2.5% of measured proteins (**Fig. 5a, Supplementary Table 2**). Gene ontology enrichment of cellular components showed an over-representation of intermediate filaments, keratin filaments and cytoskeletal proteins in the *Krt76*^-/-^/nu serum (**Fig. 5b**). Collectively, these data demonstrate that *Krt76* thymic deficiency leads to a substantial reduction in skin-specific TSA expression, providing a mechanistic link between keratin loss and the observed skin and oral mucosa autoimmune phenotype.

**Fig. 5.**
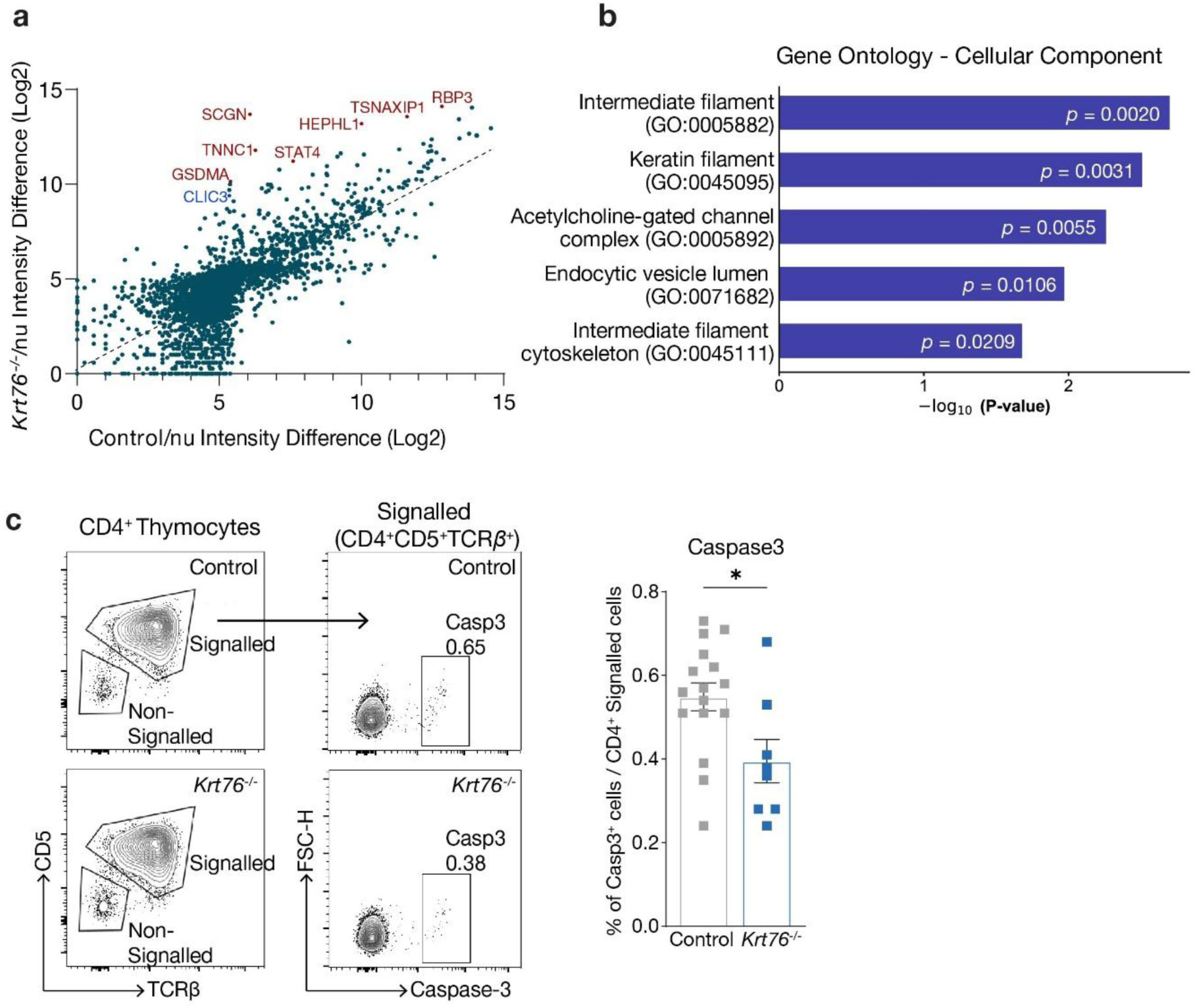
Loss of thymic *Krt76* results in autoantibody reactivity to *Krt76*-expressing TSAs and impaired negative selection. **a**, Comparison of autoantibody profiling for Control/nu and *Krt76*^-/-^/nu serum detected via filtered HuProt assays. CLIC3 signal highlighted in red. Intensity Difference (a.u.) relates to difference between positive autoantibody signal versus control IgG antibody per sample. **b**, Gene Ontology Cellular Component (2025) analysis of genes with Log2 Fold Change (*Krt76*^-/-^/nu / Control/nu) greater than 1.5 detected via filtered HuProt assays. **c**, Quantification of apoptosis of single-positive T cells in the thymus. Representative plots for flow cytometry gating of CD4^+^ signalled thymocytes (CD5^+^TCRβ^+^) and Cleaved Caspase3^+^ thymic T cell populations of Control and *Krt76*^-/-^ (4- month-old). Mean±s.e.m. t-test statistical analysis, \**p* ≥0.05. *n*=3 experiments consisting of 8-16 animals/group.

### Loss of *Krt76* impacts T cell selection

To investigate whether *Krt76* loss and the decreased expression of the *Krt76*-expressing mTECs-specific gene signature affect thymic T cell selection, we performed integrated scRNA-seq and scTCR-sequencing on thymocytes isolated from control and *Krt76*-deficient mice (**Supplementary Fig. 8a,b**). Since *Krt76* deficiency reduces the expression of skin-associated TSAs, we hypothesised that disruption of this critical aspect of the mimetic mTECs repertoire would result in altered TCR gene usage patterns. Interestingly, principal component analysis of V/J repertoire revealed differential TCR alpha chain usage in single-positive CD4+ and CD8+ T cells as well as regulatory T cell populations in *Krt76*^-/-^ mice compared to controls (**Supplementary Fig. 8c**, *p* = 0.16), suggesting subtle but meaningful shifts in the TCR repertoire composition. To explore this further we bulk sequenced the TCRβ gene from single positive CD4^+^ T cells and DP T cells from Control and *Krt76*^-/-^ mice. Bulk TCRβ-sequencing showed that *Krt76* deficiency increased CDR3 length divergence between DP and CD4^+^ SP cells (*p* = 0.057), whereas control populations remained stable (*p* = 0.34) (**Supplementary Fig. 8d,e**), suggesting differences in thymocyte selection at the negative selection phase in *Krt76*^-/-^ mice. Given that negative selection shapes the TCR repertoire primarily at the single positive (SP) stage, we reasoned that the observed SP-specific repertoire shifts in *Krt76*^-/-^ mice might result from defective central tolerance. To test this, we quantified apoptotic thymocytes undergoing negative selection. Flow cytometry revealed a significantly reduced proportion of CD4^+^ cleaved-Caspase3^+^ thymocytes in *Krt76*^-/-^ mice compared to controls (**Fig. 5c**). This reduction suggests that impaired presentation of *Krt76*-dependent TSAs enables autoreactive thymocytes to evade negative selection and escape into the periphery. These findings are consistent with a model wherein disrupted presentation of specific skin antigens by mTECs leads to reduced negative selection of autoreactive T cell clones.

Collectively, our findings reinforce the conclusion that *Krt76* is required in the murine thymus to facilitate the terminal differentiation of a subset of post-*Aire* skin antigen-expressing mimetic mTECs that are necessary to establish robust central tolerance of certain skin antigen-targeting autoreactive T cells.

## DISCUSSION

Keratins have long been viewed as guardians of epithelial integrity, with their dysfunction often linking epithelial barrier disruption to skin inflammation^3,4,34^. This study challenges that paradigm, uncovering an essential and unexpected role for KRT76 in the thymus establishing central tolerance to skin-targeting T cells. By transferring *Krt76*^-/-^ thymic lobes under the kidney capsule of nude mouse recipients, we revealed that loss of thymic *Krt76* drives a peripheral immunopathology targeting specifically the oral mucosa and skin (e.g. with autoantibodies), independent of epidermal barrier impairment^11,26^.

Mechanistically, we track this KRT76-dependent thymus defect to aberrant differentiation of a subset of mTECs that adopt keratinocyte-like characteristics, known as CorneoTECs. This mimetic mTECs III subset is thought to differentiate from a postnatal wave of TEC progenitors that first generate a proliferative mTECs I population and then AIRE-expressing mTECs II cells^35–37^. Until recently, the provision of tissue-self antigens (TSAs) for T cell negative selection was thought to be under the sole regulation of AIRE, which acts stochastically to induce a broad and unconnected set of tissue-restricted genes^22,38–41^. However, a series of evolutionarily conserved post-AIRE mimetic mTECs III subsets have now also been identified that retain mTECs identity, but also deploy lineage-defining transcription factors (TFs) to drive gene signatures and phenotypic features of differentiated peripheral cells^14,21,29,42^. Some mimetic mTECs, which include CorneoTECs, are derived from AIRE-expressing mTEC II cells^14,21,30^, in which AIRE may provide a permissive chromatin landscape on which tissue-specific TFs can act; the grainyhead-like (*Grhl*) family of TFs have been shown to regulate the CorneoTEC program^14,15^.

Interestingly, *Krt76* appears to be expressed in only a subset of both the AIRE^+^ mTECs II and AIRE^-^ mTECs III CorneoTECs populations. This may simply reflect a relatively transient expression of *Krt76* along dynamic stages of CorneoTECs differentiation. However, as *Krt76*-expressing Mimetic mTECs cells differentially express a transcriptional signature that features numerous skin and oral mucosa- restricted genes that are associated with more terminal states of differentiation (e.g. *Clic3*, *Gsta2*, *Defb6*, *Dapl1*, *Trim29*, *Aldh3b2* and *Dmkn*), this may instead indicate that KRT76 identifies a specific sub-type of CorneoTECs. Such additional granularity suggests that multiple parallel CorneoTECs programs may exist, that could each represent a specific stage of development, state of differentiation, and/or anatomical location, to cover a greater diversity of keratinocyte-associated self-antigens. These parallel CorneoTECs programs may be regulated by different combinations of keratinocyte-related TFs, that could direct distinct keratinocyte-like trajectories in the background of the stochastic, quasi-random nature of permissive chromatin accessibility set up by AIRE function in mTECs II cells^22,40,43,44^. Notably, loss of AIRE does not result in the complete absence of any mimetic cell subset, although many, but not all, are reduced in frequency and number, and express an altered TRA signature^14^. Moreover, individual thymic medullary islets derive from individual mTECs precursors^45^, supporting the idea that single mTECs precursors could be biased toward specific CorneoTEC states.

An important finding of this study is the significant ablation of the *Krt76*-associated transcriptional signature in *Krt76*-expressing CorneoTECs that lack functional KRT76 protein. This mechanistically links thymic expression of KRT76, which heretofore was considered primarily for its structural capacities, to central tolerance processes directed toward skin and oral mucosa-targeting self-reactive T cells that we demonstrate can drive autoimmune pathology in *Krt76*^-/-^ animals. Notably, we also demonstrate that loss of *Krt76* disrupts the organised differentiation of the mTECs-II to corneoTEC stages. This suggests that expression of KRT76 actively supports the development of (at least a subset of) the CorneoTECs compartment, perhaps by providing the correct internal environment and developmental feedback to induce a full set of KRT76-dependent keratinocyte-associated genes. In skin, KRT76 is recognised for regulation of Claudin-1-dependent tight junctions^25^. Such structures may thus also be important for the differentiation of KRT76-dependent CorneoTECs, with cells stalling along this trajectory if *Krt76* is absent. Alternatively, type II keratins (of which KRT76 is an example) have been shown to act as scaffold proteins that bind and anchor signalling proteins such as AKT and PKC, regulating how these molecules then participate in signal transduction^46–49^. Therefore, it is conceivable that the absence of *Krt76* may fundamentally affect signalling inputs in developing CorneoTECs that arrest their differentiation as revealed by trajectory and RNA velocity analyses, and ultimately affect the expression of TSAs in the CorneoTECs. These alternative scenarios for KRT76 function in developing CorneoTECs are active and ongoing areas of investigation. In this study, we have shown that disruption of KRT76 dismantles this antigenic landscape, resulting in a severe deficit of skin and tongue TSA expression in CorneoTECs, ultimately suggesting a failure of negative selection of skin-specific autoreactive T cells. The specificity of the ensuing autoimmune response – directed at skin and oral mucosa – closely mirrors this antigenic gap, tightly linking cellular mimicry with organ-selective tolerance outcomes.

Finally, our study prompts a broader re-evaluation of skin immunopathologies induced by keratin gene mutations. Until now, these inflammatory manifestations were principally thought to be a consequence of the breakdown of epithelial barrier integrity, allowing greater opportunity for epidermal infections^3,4,50^. While clearly a critical component, our data invite a further consideration: that keratin mutations may also disrupt thymic keratinocyte-like mimetic cell differentiation, resulting in aberrant skin-associated self-antigen expression, and a failure of central tolerance to skin (and oral mucosa)-specific T cells. As a consequence, inflammatory responses in the skin of affected individuals may also consist of an autoimmune component that would not, for example, resolve as a result of antibiotic regimens, warranting re-evaluation of current treatment strategies.

Looking ahead, decoding the full breadth of these thymic mimetic programs – across organs and disease states – remains a thrilling challenge poised to better understand thymus biology.

---

**Supplementary Fig. 1.**
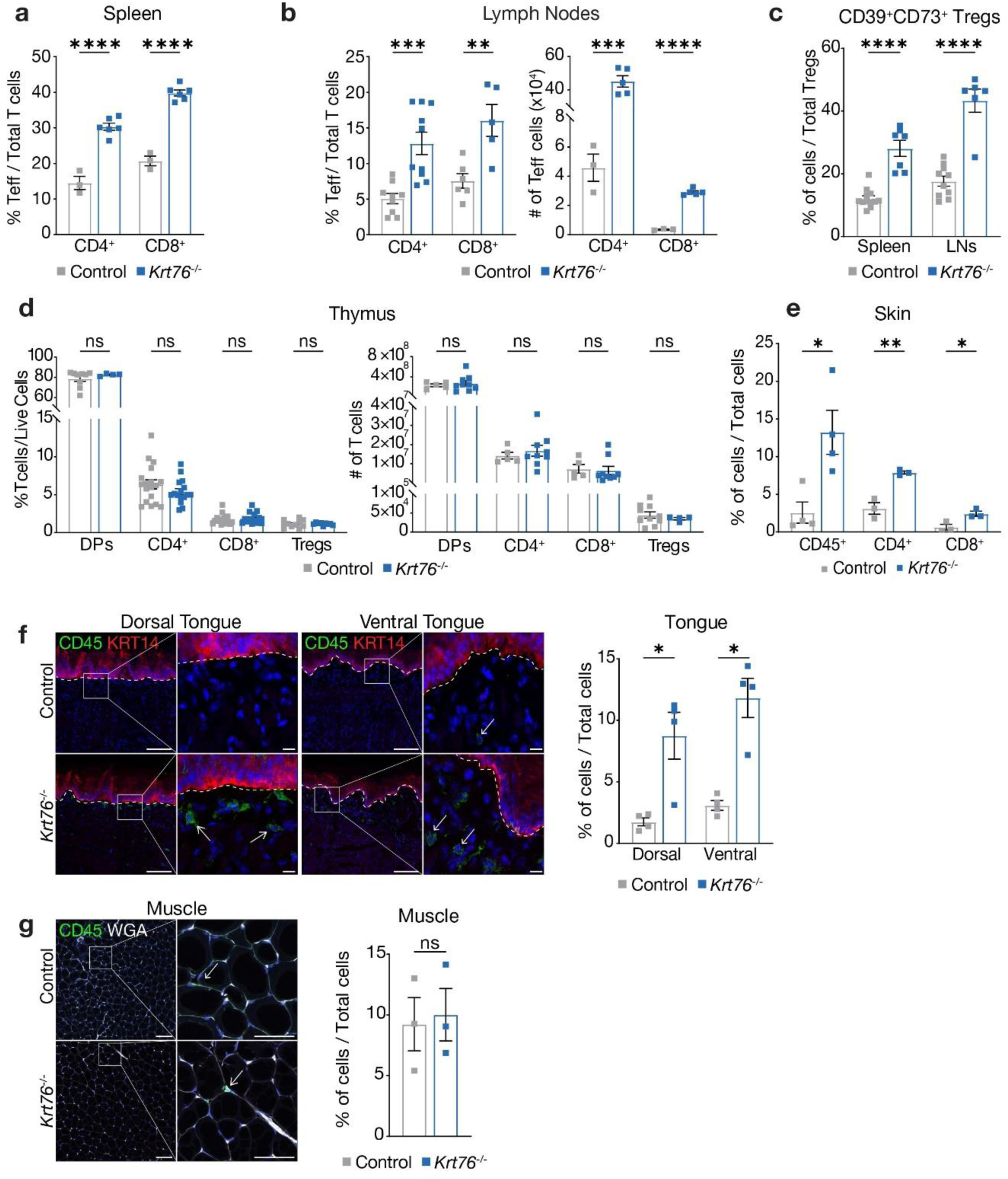
*Krt76*^-/-^ mice show an immune imbalance in the periphery. **a**, Flow cytometry analysis of P28 splenic CD4^+^ and CD8^+^ effector T cells for Control and *Krt76*^-/-^ mice. **b,** Flow cytometry analysis of P28 lymph node CD4^+^ and CD8^+^ effector T cells for Control and *Krt76*^-/-^ mice with their respective absolute numbers. **c**, Flow cytometry analysis of spleen and lymph node CD39^+^CD73^+^ Tregs (CD4^+^CD25^+^FoxP3^+^) for P28 Control and *Krt76*^-/-^ mice. **d**, Flow cytometry analysis of P28 thymocyte populations Double Positive (DPs; CD4^+^CD8^+^), CD4^+^ Single Positive (CD4^+^CD8^-^), CD8^+^ Single Positive (CD4^-^CD8^+^) from live cells, and T regulatory cells (Tregs; CD25^+^FoxP3^+^) from CD4+ Single Positive for Control and *Krt76*^-/-^ mice with their respective absolute numbers. **e**, Proportion of infiltrating CD45^+^/CD4^+^/CD8^+^ cells/mm^2^ in skin dermal regions versus total cells in Control and *Krt76*^-/-^ mice back skin. **f**, Representative immunofluorescence staining of P28 dorsal and ventral tongue using anti-KRT14 (red), anti-CD45 (green, arrows) and DAPI (nuclear counterstain), with quantification of infiltrating CD45^+^ immune cells/mm^2^ in dorsal and ventral tongue of Control and *Krt76*^-/-^ mice versus DAPI^+^ cells. **g**, Representative immunofluorescence staining of P28 femur skeletal muscle of Control and *Krt76*^-/-^ mice using anti-CD45 (green, arrows), DAPI (nuclear counterstain) and WGA (white, membrane counterstain), with quantification of infiltrating CD45^+^ immune cells/mm^2^ in muscle of Control and *Krt76*^-/-^ mice versus DAPI^+^ cells. Mean±s.e.m. t-test statistical analysis, *ns=*non-significant, \**p*≤0.05, \*\**p*≤0.01, \*\*\**p*≤0.001, \*\*\*\**p*≤0.0001. *n*=2-5 experiments consisting of 3-18 animals/group. Scale bars: 10μm, 50μm or 100μm (f, g).

**Supplementary Fig. 2.**
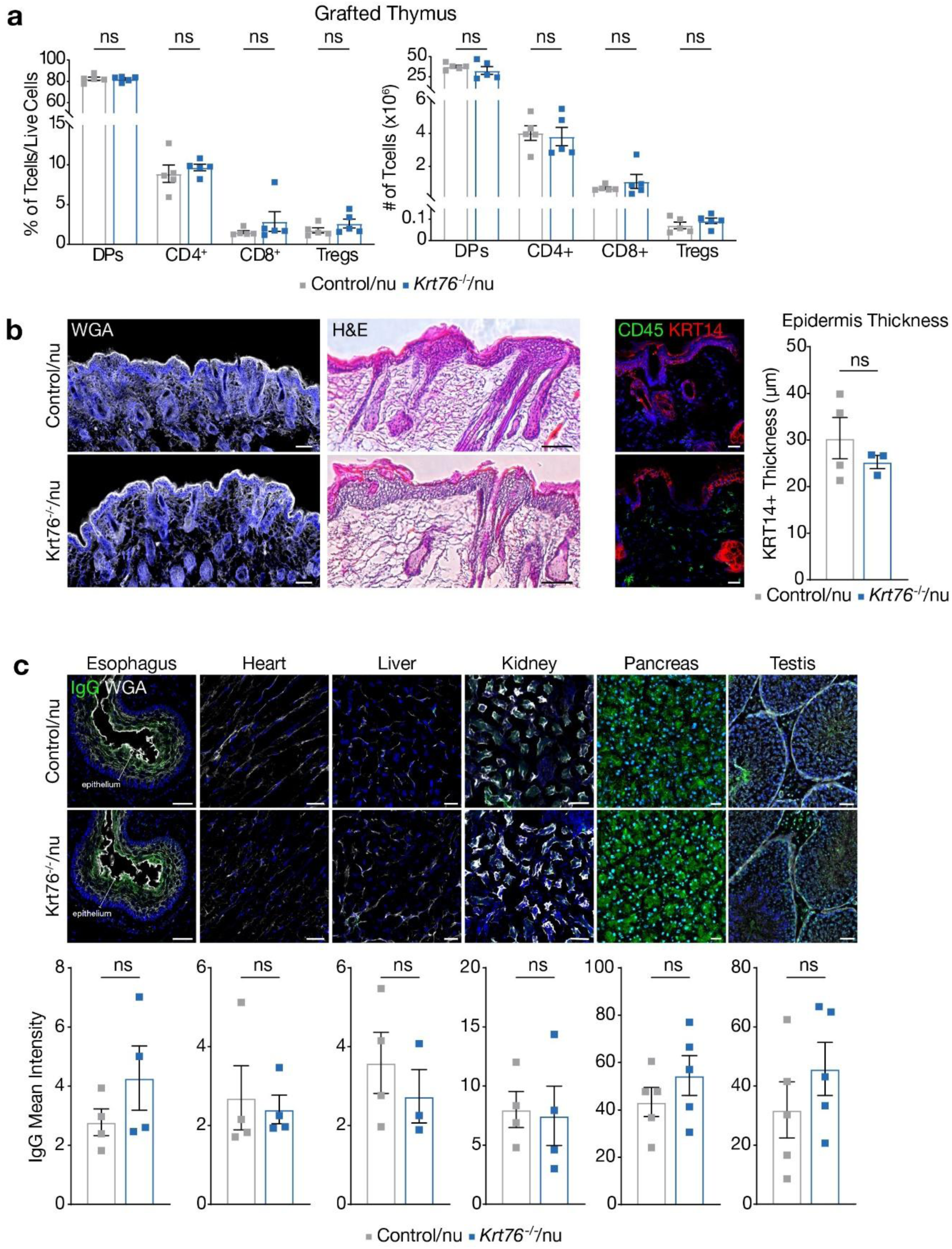
Loss of Thymic *Krt76* drives tissue-specific autoimmune response. **a**, Flow cytometry of Control/nu and *Krt76*^-/-^/nu thymocyte populations: Double Positive (DPs; CD4^+^CD8^+^), CD4^+^ Single Positive (CD4^+^CD8^-^), CD8^+^ Single Positive (CD4^-^CD8^+^) from live cells, and T regulatory cells (Tregs; CD25^+^FoxP3^+^) from CD4^+^ T cells, 10 weeks post transplantation. **b**, Immunofluorescence membrane counterstain (WGA, white), hematoxylin-eosin (H&E) stained sections, and staining and quantification of KRT14^+^ (red) epidermis thickness in back skin of grafted Control/nu and *Krt76^-^*^/-^/nu mice. **c**, Immunofluorescence staining and quantification of IgG autoantibodies in sera of Control/nu and *Krt76*^-/-^/nu mice detected by staining *Rag1*^-/-^ mice oesophagus, heart, liver, kidney, pancreas and testis sections, and measuring the mean fluorescent intensity. Scale bars: 100μm or 1mm (b), 25μm (c).

**Supplementary Fig. 3.**
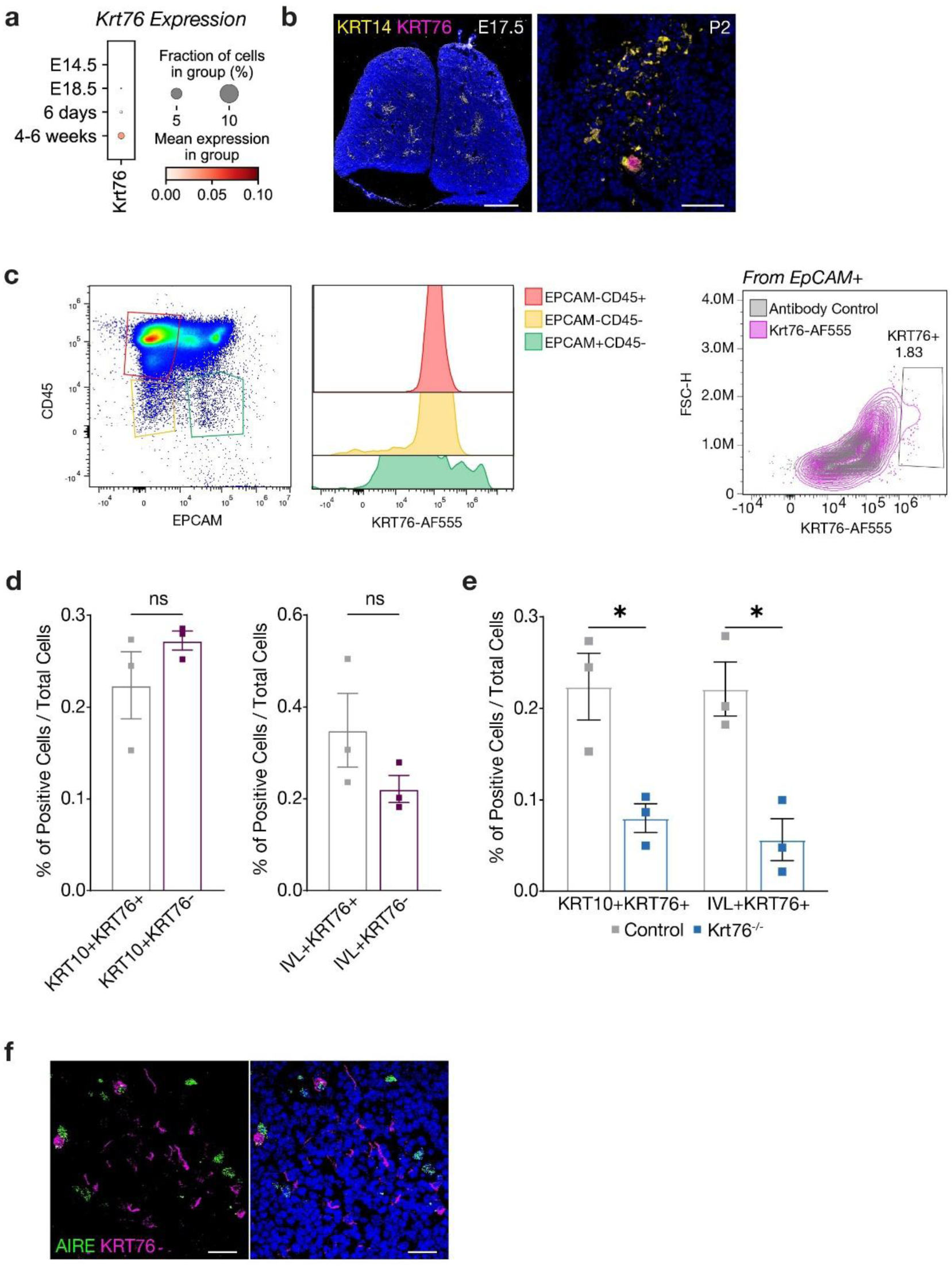
KRT76 Expression in the Thymus. **a**, Dotplot of scRNA-seq thymic epithelial cell subpopulations in mouse datasets with *Krt76* expression across different ages (GEO: E-MTAB-8581^28^). **b**, Immunofluorescence staining of Embryonic Day 17.5 (E17.5) and Postnatal Day 2 (P2) C57/BL6 mice thymus using anti-KRT14 (yellow), anti-KRT76 (magenta) and DAPI (nuclear counterstain). **c**, Representative flow cytometry dot plot showing expression of KRT76 in CD45-EpCAM^+^ (green) populations and KRT76^+^ expression (magenta) versus AF555 antibody control (grey) in EpCAM^+^ population from adult C57/BL6 mice. **d**, Immunofluorescence quantification of double-positive populations (KRT10^+^KRT76^+^ or IVL^+^KRT76^+^) versus KRT76^-^ populations in DAPI^+^ total cells within medullary areas (KRT14^+^). **e**, Immunofluorescence quantification of double-positive populations (KRT10^+^KRT76^+^ or IVL^+^KRT76^+^) in Control or *Krt76*^-/-^ mice thymus in DAPI^+^ total cells within medullary areas (KRT14^+^). **f**, Immunofluorescence staining of adult C57/BL6 mice thymus using anti-AIRE (green), anti-KRT76 (magenta) and DAPI (nuclear counterstain). For IHC quantification, n≥3 sections/3-4 animals/group. Mean±s.e.m. t-test statistical analysis, *ns=*non-significant, \**p*≤0.05. Scale bars: 25μm (f), 50μm (b), 500μm (b).

**Supplementary Fig. 4.**
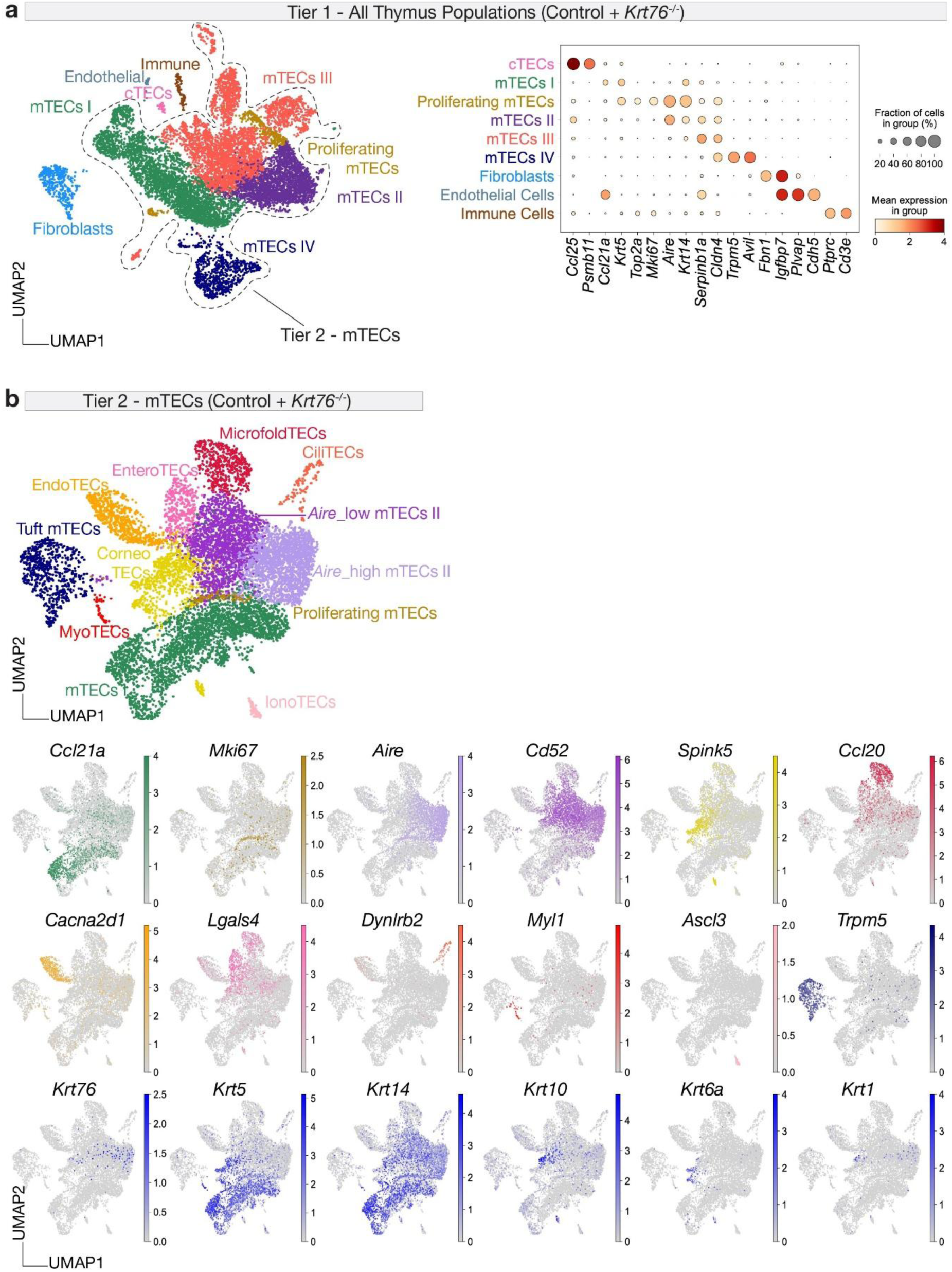
Loss of *Krt76* impacts thymic cell differentiation. **a,** scRNA-seq UMAP annotation and dotplot summary of representative markers for Tier1 clustering of all thymus populations identified following integration of Control and *Krt76*^-/-^ samples; total cell number post-QC filtering (Control=4,271 cells; *Krt76^-/-^*=7,343 cells). Dotted line delineates Tier 2 mTECs clusters selected for further downstream analysis. **b**, scRNA-seq UMAP visualisation of Tier 2 subclustering of integrated Control and *Krt76^-/-^*mTECs subpopulations, total cell number post-QC filtering (Control=3,650 cells; *Krt76^-/-^*=5,909 cells) with UMAP visualisation of each mTECs clusters’ representative markers in Tier2 and *Krt76, Krt5, Krt14, Krt10, Krt6a* and *Krt1* expression across mTECs subsets. *n*=2 scRNA-seq experimental repeats.

**Supplementary Fig. 5.**
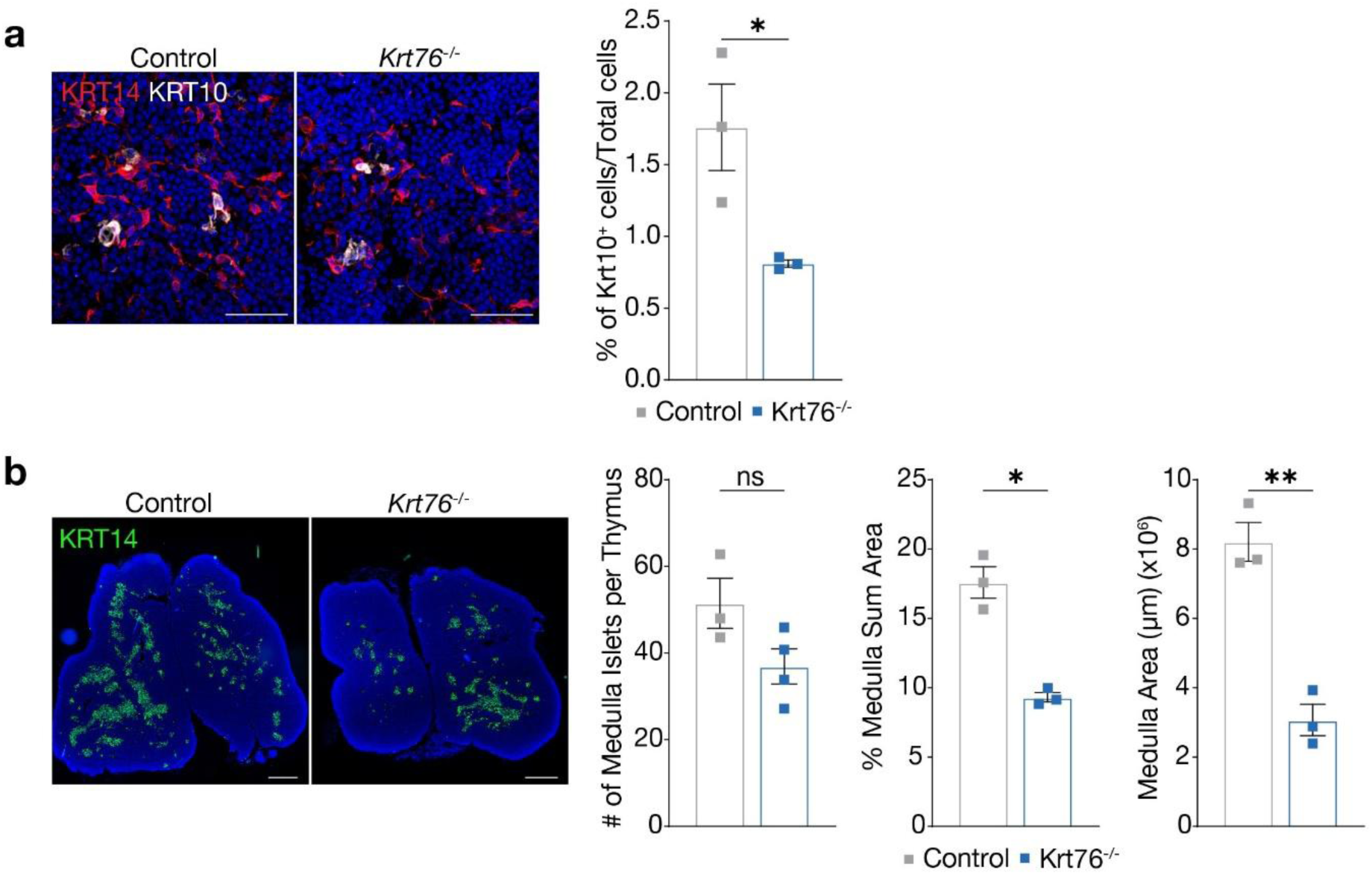
Loss of *Krt76* impacts thymic cell differentiation. **a,** Immunofluorescence staining of Control and *Krt76*^-/-^ mice thymus with anti-KRT14 (red), anti-KRT10 (white) and DAPI (nuclear counterstain), with quantification of KRT10^+^ cells versus DAPI^+^ total cells within medullary areas (KRT14^+^). **b**, Immunofluorescence quantification of medullary islets (KRT14^+^) per thymus, proportion of the average medullary areas (KRT14^+^) relative to each thymus area and total medullary area (KRT14^+^) from *n*=11-18 sequential sections of Control and *Krt76^-/-^* mice with anti-KRT14 (green) and DAPI (nuclear counterstain). For IHC quantification, n≥6 sections/3 animals/group. Representative of *n*=3 experiments consisting of 3-4 animals/group. Mean±s.e.m. t-test statistical analysis, *ns=*non-significant. \**p*≤0.05, \*\**p*≤0.01. Scale bars: 100μm (a), 1mm (c).

**Supplementary Fig. 6.**
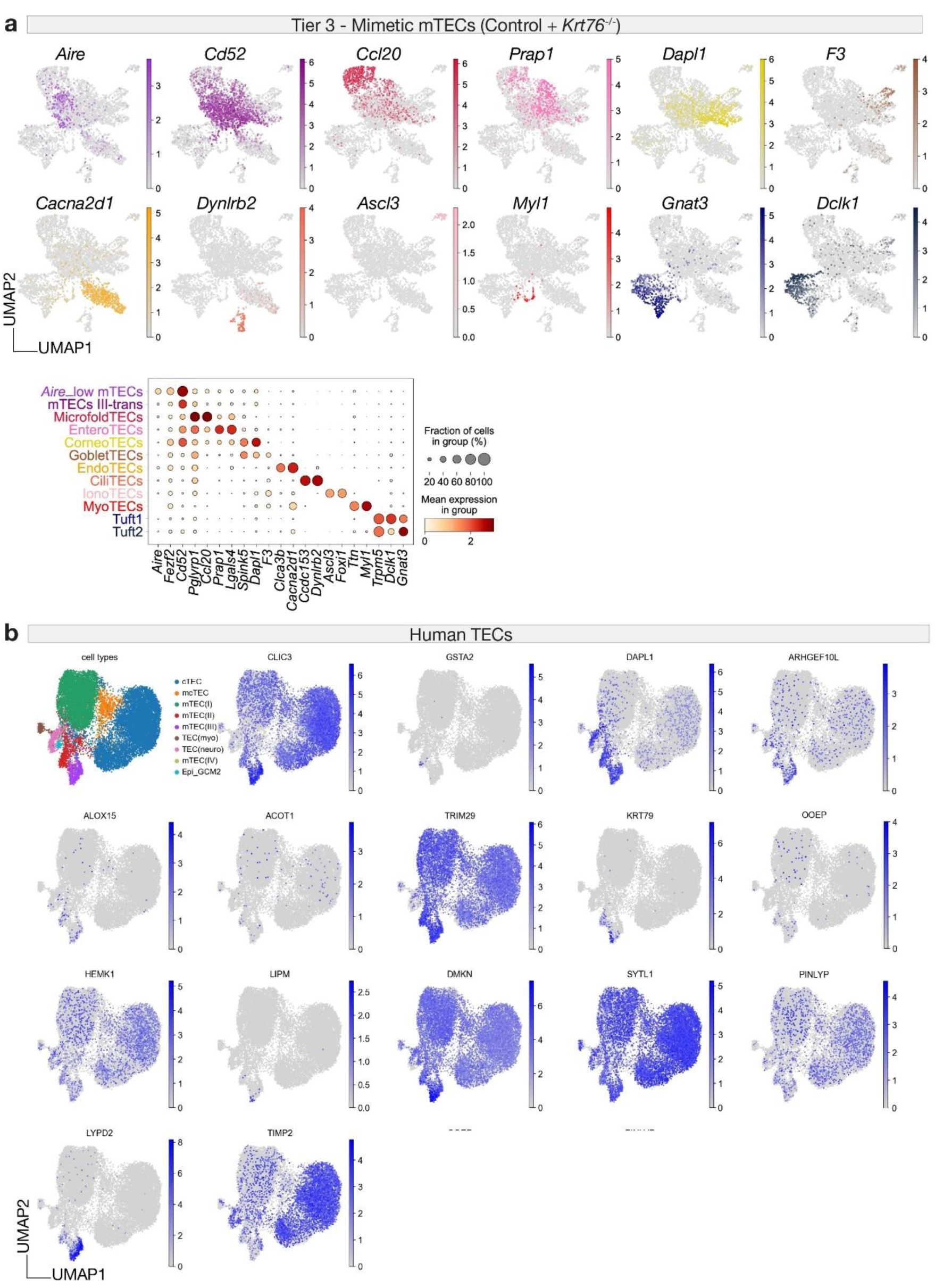
Expression of *Krt76*-TSA signature in mouse and human mTECs. **a**, UMAP and dotplot annotation of Mimetic mTECs subpopulations based on representative markers in Control and *Krt76*^-/-^ sample. **b**, Dotplot with expression of Krt76-specific TSAs in human TECs (GEO: E-MTAB-8581)^28^.

**Supplementary Fig. 7.**
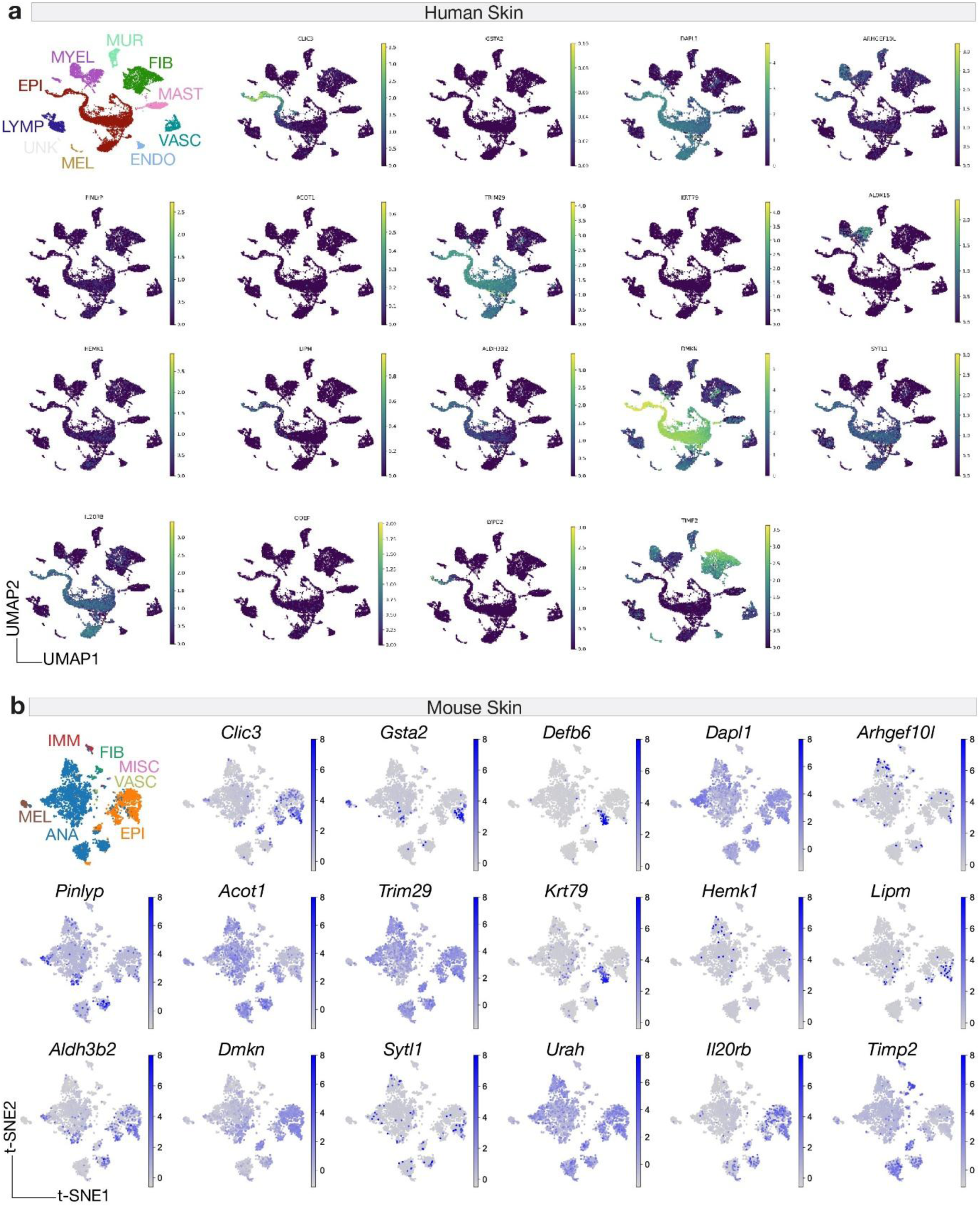
TSAs found in *Krt76*-expressing mTECs are expressed in human and mouse skin (supporting Figure 4b). **a,** Expression of *Krt76*-TSA signature in healthy adult human skin scRNA-seq dataset (GEO: GSE241132)^51^, analysed in Skin explorer^52^. UMAPs showing the expression of *CLIC3, GSTA2, DAPL1, ARHGEF10L, PINLYP, ALOX15, ACOT1, TRIM29, KRT79, HEMK1, LIPM, ALDH3B2, DMKN, SYTL1, IL20RB, OOEP, LYPD2* and *TIMP2*. EPI: Epidermis keratinocytes; FIB: Fibroblast-like cells; MYEL: myeloid Cells; LYMP: lymphocytes; MAST: mast cells; ENDO: endothelial cells; MEL: melanocytes; MISC: Miscellaneous; VASC: Vascular cells; UNK: unknown. **b**, t-SNE plot visualisation of *Krt76*-expressing mTECs’ TSAs in a published adult wild-type mouse skin dataset (GEO: GSE129218)^31^. ANA: Anagen hair follicle keratinocytes; EPI: Permanent epidermis keratinocytes; FIB: Fibroblast-like cells; IMM: Immune Cells; MEL: melanocytes; MISC: Miscellaneous; VASC: Vascular cells.

**Supplementary Fig. 8.**
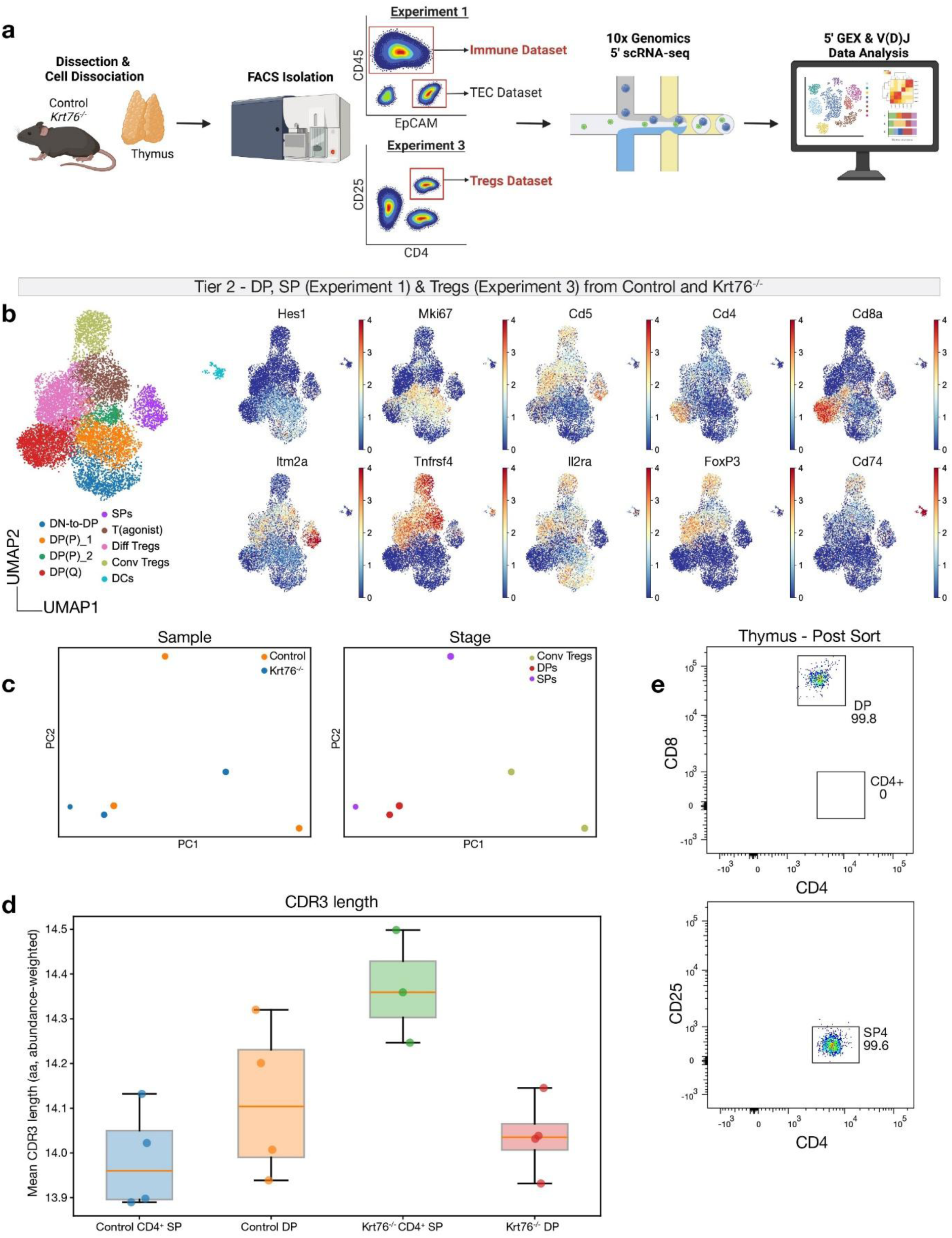
Loss of *Krt76* impacts T cell negative selection. **a**, scRNA-seq experimental workflow, thymi were collected from Control and *Krt76*^-/-^ 4-6 weeks mice (*n*=4 animals/genotype), FACS sorting was used to collect thymic immune populations, either CD45^+^ cells (Experiment 1) or CD4^+^CD25^+^ cells (Experiment 3). **b**, UMAP annotation of merged Double-Positive (DP), Single-Positive (SP) from Experiment 1 and Regulatory T cells (Tregs) subtypes from Experiment 3 with representative markers for each subpopulation. **c**, Principal component analysis (PCA) of TCRα V/J feature space pseudobulk for Control and *Krt76*^-/-^ of DP (merged DP(P)_1, DP(P)_2, DP(Q)), SP and Conventional Tregs (Conv Tregs) subsets using Dandelion package^53^. Each point represents a cell type pseudobulk. **d**, UMI-abundance-weighted mean CDR3 length from TCRβ bulk sequencing of Control CD4^+^ SP, Control DP, *Krt76*^-/-^ CD4^+^ SP and *Krt76*^-/-^ DP samples, each point represents a sample replicate. *Krt76*^-/-^ CD4^+^ SP versus *Krt76*^-/-^ DP CDR3 length *p* = 0.057, Control CD4^+^ SP versus Control DP CDR3 length *p* = 0.34 (two-sided Mann-Whitney U test). **e**, Representative dot plots of thymic double positive (DP) and CD4^+^ SP (SP4) T cell populations post-sort purity check for TCRβ sequencing. Image in **a** created with Biorender.com.

**Supplementary Fig. 9.**
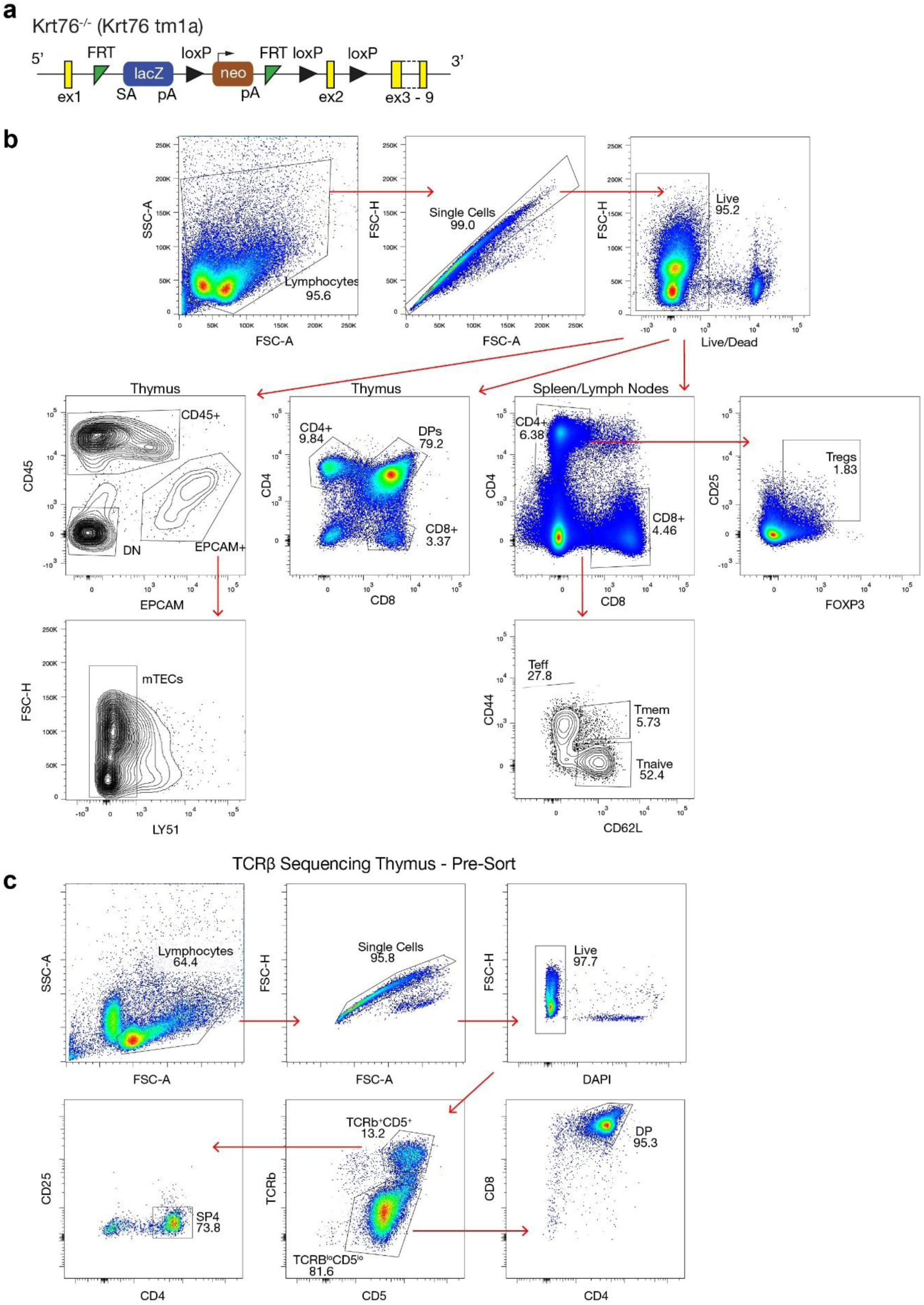
**a**, *Krt76* knockout strategy (Krt76 tm1a). **b,** Flow cytometry plots of the gating strategy used to analyse live TECs (CD45^-^EPCAM^+^) and medullary TECs (CD45^-^EPCAM^+^LY51^-^); thymus, spleen and lymph nodes (LNs) CD4^+^ and CD8^+^ populations, including Tregs from CD4^+^ T cells (CD25^+^FOXP3^+^) and respective T cell subpopulations (CD44 and CD62L). **c**, Gating strategy for thymic T cell populations pre-sort for TCRβ sequencing of signalled TCRβ^hi^CD5^hi^CD4^+^CD25^-^ single positive CD4^+^ T cells (SP4) and non-signalled TCRβ^lo^CD5^lo^CD4^+^CD8^+^ double positive T cells (DP).

**Supplementary Table 1. Differential expression analysis of tissue-restricted antigens in *Krt76*⁺ Mimetic mTECs.** The table lists genes preferentially expressed in Krt76⁺ mTECs identified by scRNA-seq differential expression analysis using the *sc.tl.rank_genes_groups* with the statistical method set to ‘Wilcoxon’ and filtering for outputs with a fold-change≥1.5 using *sc.tl.filter_rank_genes_groups* to ensure statistical significance.

**Supplementary Table 2. IgG autoantibody protein specificity analysis of Control/nu and *Krt76*^-/-^ /nu serum using the HuProt array.** Table outlining analysis steps to identify Log2 Intensity Difference and Log2 fold change differences between *Krt76*^-/-^/nu and Control/nu serum against protein hits from the HuProt Human Proteome Microarray v4.

## ACKNOWLEDGEMENTS

We thank past and present members of the Sequeira group for their feedback and support, special thanks to Yufan Yao; the staff of the animal facility at QMUL, KCL and The Francis Crick Institute, including Matteo Battilocchi and Ian Goodison; Karl Annusver for his advice on the RNA velocity analysis; the staff of the QMUL Genome Centre, Flow facility, Microscopy and Phenotypic Screening Facility. *Rag1*^-/-^ mice (*C57BL/6J-Rag1em10Lutzy/J*) were kindly donated by Dominique Bonnet. We are grateful to David Kelsell and Kif Liakath-Ali for insightful discussions and constructive feedback on the manuscript.

## AUTHOR CONTRIBUTIONS

I.S. conceived the project. M.G., P.B., D.P. and I.S. designed the project. M.G. performed most of the experiments and analysis. P.B. performed the renal capsule surgery. R.R. and S.C. helped with graft harvesting. R.R. performed the *in vitro* cultures, RNA extractions and qPCR experiments. M.G., I.T. and I.S. performed scRNA-seq library preparation. M.G. performed scRNA-seq analyses, with support from M.E.. M.S., N.Y. and K.T. performed the V(D)J and TCRβ analyses. M.G., I.T., R.R., M.C., M.V.R., D.P., I.Sa. and M.T. performed and analysed histology and immunofluorescent experiments. N.S. and I.Sa. provided support on flow cytometry experiments and analysis. F.W., P.B., M.G. and I.S. acquired funding for the project. M.G. D.J.P. and I.S. wrote the manuscript with input from all authors. All authors read and approved the final manuscript.

## MATERIAL AND METHODS

### Human samples

Human thymic tissue used for immunofluorescence staining of Hassall’s Corpuscles and to generate thymic epithelial stem cell cultures was obtained from patients undergoing cardiothoracic surgery at the Great Ormond Street Hospital (REC No 15/YH/0334 and 07/Q0508/43-06-MI-13). Patients or legally authorised representatives gave written informed consent.

### Animals

The *Foxn1^nu^* (*CAnN.Cg-Foxn1nu/Cr*) mice were purchased from Charles River Laboratories. Krt76^-/-^ mice (*Krt76^tmla(KOMP)Wtsi^*) (**Supplementary Fig. 9a**) were obtained from the Wellcome Trust Sanger Institute Mouse Genetics Programme, as part of the International Mouse Phenotype Consortium^54^. The Krt76^-/-^ strain was designed by selecting the 5’-most critical exons which disrupt at least 50% of the protein-coding sequence. The *Krt76* gene is interrupted due to the splicing of the LacZ trapping element at the *Krt76* exon. Although the protein is disrupted, exon1 and *LacZ* reporter gene between the first and second exons allows tracking of *Krt76* expression. Crossing of heterozygous (Krt76^+/-^) mice produced viable knockout Krt76^-/-^ mice, as well as littermates Krt76^+/-^ and wild type (Krt76^+/+^). Krt76^+/-^ shows no phenotype compared to Krt76^+/+^ ^11^ and were used as Control. The Rag1^-/-^ mice (*C57BL/6J-Rag1em10Lutzy/J*) were provided by D. Bonnet and P. Bonfanti. Mice were maintained on the C57Bl/6 N genetic background and kept at the Francis Crick Institute, King’s College London and Queen Mary University of London animal facilities. All handling and procedures were conducted under the guidelines of the UK Home Office and performed under the terms of UK Home Office Project Licences (PP5184214, PP9619702 and PPL70/19296).

### Mouse genotyping

Mice genotyping was performed via polymerase chain reaction (PCR) following DNA extraction using DNA isolation buffer (100mM Tris, 50mM KCl, 2mM MgCl2, 0.1mg/ml Gelatine, 0.45% NP40, 0.45% Tween, 20mg/ml Proteinase K (New England BioLabs, P8107S)). Tissue was incubated in DNA extraction buffer at 55°C for 2-3 hours or overnight, followed by 10min 95°C enzymatic inactivation. the KAPA Mouse Genotyping Kit (KAPA Biosystems, KK7301) was used to prepare the PCR master mix as per manufacturer instructions with 10μM forward (GAGCAAGAGGATTTCCTGAGAACTC) and reverse primers (GCTTAAAAATAGCAGCAACTGGAACA), and 10ng of template DNA.

### Renal Capsule Transplantation of Embryonic Thymi

E15.5 thymi from Krt76^-/-^ and Control mice were harvested and collected in RPMI 1640 Medium (ThermoFisher Scientific, 11875093) + 2% heat-inactivated FBS (ThermoFisher Scientific, 10-500-064). Embryo’s genotype was determined using the method outlined above. The kidney of athymic recipient 2-3 months old *CAnN.Cg-Foxn1nu/Cr* (Nude, nu) mice (Charles River Laboratories) was exposed via a small dorsolateral cut. The kidney’s capsule was incised and a single E15.5 thymus was transplanted into a single recipient nude mouse renal subcapsular, similarly to previous reports^55^. Overall, 6 transplants/genotype were performed at each experiment. The mice were maintained under observation for 10 weeks in pathogen-free conditions. After 10 weeks, animals were sacrificed and spleen, submandibular and axillary lymph nodes, blood and back skin were harvested from all grafted mice. Spleen weight was measured. Spleen and lymph nodes were collected for flow cytometry. Skin was collected for immunostainings following the method outlined below. Following 30-60min at RT, blood was centrifuged 10min 1000g to collect serum and stored at -80°C for future analysis. Workflow schematic outlined in Fig. 1d.

### Peripheral Tissue Preparation for Flow Cytometry

Spleen was harvested from P28 Control and Krt76^-/-^ mice, crushed through a 40μm cell strained (Falcon, 352340) and rinsed with chilled PBS. Cells were treated with 1x red blood cell (RBC) lysis buffer (eBioscience, 00-4333-57) for 5min at RT. All cells were collected by centrifugation and strained through a 40μm cell strainer. Two axillary and two submandibular lymph nodes were harvested from P28 Control and Krt76^-/-^ mice, they were merged and crushed through a 40μm cell strained and rinsed with chilled PBS. Thymus from 4-6 weeks old Control and Krt76^-/-^ mice were crushed in 0.5mL chilled PBS using a pestle and strained through a 40μm cell strained to remove thymocytes. The remaining stromal tissue was finely sliced and digested enzymatically using 0.05 % [w/v] of Liberase (Roche, 5401119001), 100 U/ml DNase I (Sigma-Aldrich, 10104159001) in RMPI 1640 Medium (ThermoFisher Scientific, 11875093) for 30-40min at 37°C at 100rpm. Cells were mechanically disrupted using a 21G needle to break up aggregates and filtered through at 100µm filter for single cell suspension and depleted for CD45-expressing cells via immunomagnetic separation (EasySep Mouse TIL (CD45) Positive Selection Kit, StemCell, 100-0350) and used for mTECs populations staining. 4-months old Control and Krt76^-/-^ mice were crushed in 0.5mL chilled PBS using a pestle and strained through a 40μm cell strained to collect thymocytes for Cleaved Caspase3^+^ staining.

### Differentiation of human thymic epithelial cultures

Human thymic epithelial cultures were prepared and differentiated as previously described^29^. Briefly, Thymic PolyKRT stem cells were sorted using FACS from single live (Zombie Aqua Fixable Viability Kit, Biolegend, 423102) BCAM+ cells (BCAM BV421 (BD, 748007)) and seeded in culture conditions favouring their expansion from donor thymi^29^. 90,000 PolyKRT cells (800/mm^2^) were plated on a membrane insert for well plates (Greiner Bio) and cultured for two days in complete FAD (Expansion media) composed of 3:1 mixture of DMEM 1X (Gibco, 41965039) and F-12 Nut Mix (Gibco, 21765029), 10% FBS (Sigma-Aldrich, F2442), 1% penicillin and streptomycin (100X, Sigma-Aldrich, Cat: p4333-100), Hydrocortisone (0.4 mg/ml, Calbiochem, 386698), Cholera Toxin (10^-10^M, Sigma-Aldrich, Cat: 80052), Triodothyronine (T3) (2×10^-9^M Sigma-Aldrich, Cat: 80052) and Insulin (5 mg/ml, Sigma-Aldrich, 91077C). All reagents were filtered through a 0.22mm strainer. From day 3, expansion media was removed, and differentiation media was added to the plated cells (PneumaCult, STEMCELL Technologies) and changed every other day for 25-30 days. Cells harvested for immunofluorescence were washed three times with PBS and fixed with 4% PFA for 10min, after which the cells were washed again three times in PBS and used immediately or stored at 4°C. Meanwhile, cells used for gene expression analysis were collected in ReliaPrep kit (Promega, Z6011) as described below and stored at -80°C until RNA extraction.

### RNA extraction

Cells were lysed using the Lysis buffer + Thyoglycerol buffer from ReliaPrep Kit. 100% isopropanol was added to the lysate, followed by RNA Wash Solution steps. The lysate was precipitated into a final collection elution tube and re-suspended in nuclease free water (Qiagen, 129117). RNA concentration was calculated using NanoDrop1000 (ThermoFisher Scientific) and converted to cDNA using the GoScript Reverse Transcriptase kit (Promega, A5000). The output cDNA concentration was adjusted to 10ng/ml and used in qPCR using the PCR master mix PrecisionPLUS-R, (Primerdesign Ltd, PPLUS-LR) with low-ROX and Taqman qPCR probes (Integrated DNA Technology). The reaction was loaded onto MicroAmp Fast Optical 96 well Reaction Plates (Applied Biosystems, 4346907) and read using the QuantStudio 3 Real-Time PCR System (Applied Biosystems). Their expression was then normalised to HPRT housekeeping gene to calculate the relative gene expression.

### Flow Cytometry

Single cells were resuspended to 1×10^6^ cells/ml and labelled with LIVE/DEAD Fixable Viability Dye (Invitrogen, L34962) to discriminate from dead cells and blocked for nonspecific binding with anti-CD16/CD32 Fc Block (BDPharmigen, 553141). Cells were labelled for 30min at 4°C with the following cell surface antibodies: CD8 BV605 (Biolegend, 100743), CD4 BUV395 (BDBiosciences, 740208), CD44 AlexaFluor700 (Biolegend, 103026), CD62L PerCP Cy5.5 (eBiosciences, 45-0621-82), CD25 (eBiosciences, 47-0251-82), CD39 PE-eFluor610 (eBiosciences, 61-0391-80), CD73 FITC (Biolegend, 127220), CD45 BV510 (Biolegend, 103138), EpCAM APC Cy7 (Biolegend, 118218), MHC-II AF700 (Biolegend, 107622), UEA1 Biotin (Vector Laboratories, B-1065), Streptavidin BV605 (Biolegend, 405229), AIRE FITC (eBioscience, 53-5934-82), CD104 PerCP Cy5.5 (Biolegend, 123614). Intracellular staining against FoxP3 PE-Cy7 (eBioscience, 25-5773-82), KRT76 (Atlas Antibodies, HPA019656) and Cleaved Caspase3 PE (Cell Signalling, 12768S) or Cleaved Caspase3 FITC (BD Biosciences, 570410) was performed after cells fixation with FoxP3 Fixation Buffer and Permeabilisation Buffer (eBioscience, 00-5523-00) according to manufacturer’s protocol. KRT76 expression was detected via secondary antibody staining against anti-rabbit AF555 (Life Technologies, A31572). Labelled cells were analysed on BD LSRII cell analyser (v9.0) or Cytek Aurora (v3.3.0) and data analysed using FlowJo software (v10), gating strategy outlined in Supplementary Fig 9.

### Tissue Collection and Embedding for Frozen Sections

Thymus, back skin, dorsal and ventral tongue, and skeletal femur muscle tissue were collected for frozen sections and fixed using 4% paraformaldehyde (PFA, Sigma-Aldrich, P6148)/PBS for 20min at 4°C under agitation. Following PBS wash, tissues were snap frozen in optimal cutting temperature (OCT) buffer (VWR International, 00411243) in Leica Histomolds (Leica Microsystems, Wetzlar, Germany) and stored at -20°C or used immediately for sectioning. 5-10μm thick sections were cut for thymic tissue, and all other tissues were cut at 10-20μm using Cryostat Bright (Leica Biosystems). Sections were attached to SuperFrost Microscope Slides (Epredia, 10149870) and stored at -80°C.

### Immunofluorescence staining

Frozen sections were briefly post-fixed in 2% PFA/PBS. Tissue was permeabilised using 0.5% Triton X100 (Sigma, X100-100ML) and incubated at RT 60min with blocking buffer 10% FBS, 3% BSA, 0.05% Triton X100 to avoid nonspecific binding followed by primary antibody staining incubation overnight at 4°C using the following primary antibodies against: KRT76 (Atlas Antibodies, HPA019656), KRT14 (Covance, 906001; Biolegend, 906004), KRT10 (Covance, PRB-159P-100), Involucrin (in-house, clone SY7^56^), CD45 (BioLegend, 103102), CD4 FITC (eBioscience, 11-0043-82), CD8α FITC (eBiosciences, 11-0081-82). The following secondary antibodies were incubated for 60min at RT: rat AF488 (Invitrogen, A21208), chicken AF555 (Invitrogen, A32932), chicken AF647 (Life Technologies, A21449), rabbit AF647 (Life Technologies, A31573), mouse AF555 (Invitrogen, A31570), DAPI used for nuclear counterstain (Biolegend, 422801) and WGA used for membrane counterstain (Invitrogen, W32466). Slides were mounted using ProLong antifade medium (Invitrogen, P36930). Images were acquired using a Nikon A1 Upright Confocal or Zeiss 880 LSM microscopes on the Carl Zeiss ZEN 3.5 software (blue edition), or PhenoImager Fusion (Akoya Biosciences, v2.4.0). A minimum of three sections/animal were used for quantification analysis. The imaged region of interest (ROI) for all tissues was selected based only on DAPI positive staining; for skin sections, this encompassed the full dermal depth. For analysis of KRT10^+^ cells, images were acquired at high resolution using a 20x or 40x objective, as a tile scan of the whole thymus section or within a region of KRT14^+^ staining (medullary area). For analysis of KRT14+ medullary areas, whole thymus sections were imaged using the PhenoImager Fusion (Akoya Bioscience, v2.4.0).

### H&E Staining

10µm frozen tissue section slides were air-dried for 5 minutes, fixed 4% PFA for 10 mins and washed with running tap water for 3 minutes. Next, slides were placed in haematoxylin for 4 mins, followed by 5 mins wash. Slides were dipped in 1% hydrochloric acid differentiation solution for 30 seconds and washed again. The final staining step comprised 3 mins in eosin, followed by a wash. Next, the slides went through dehydration steps with increasing concentrations of ethanol (70%-90%-100%-100%) for 30 seconds each and twice with xylene for 2 mins for clearing. The slides were mounted with DPX mounting medium and allowed to dry overnight. Images were acquired using an EVOS™ M5000 Imaging System (Thermo Fisher Scientific).

### Detection of Autoantibodies

Rag1^-/-^ back skin, dorsal tongue, squamous stomach, muscle, oesophagus, liver, heart, pancreas, kidney and testis were sectioned at 20μm thickness and incubated with 1/100 diluted sera from Control/nu and Krt76^-/-^/nu mice in blocking buffer (10% FBS, 3% BSA, 0.05% Triton X100), followed by incubation with goat anti-mouse AF488 (Invitrogen, A11029), DAPI was used for nuclear counterstain (Biolegend, 422801) and WGA used for membrane counterstain (Invitrogen, W32466). Images were colleced using Zeiss 880 LSM microscopes on the Carl Zeiss ZEN 3.5 software (blue edition) or PhenoImager Fusion (Akoya Bioscience, v2.4.0). A minimum of three sections/animal were used for quantification analysis. The imaged region of interest (ROI) for all tissues was selected based only on DAPI positive staining; for skin sections, this encompassed the full dermal depth.

### Imaging quantification

For quantification of medullary areas, ROIs delineated KRT14+ medulla islets and the whole thymus tissue to extract the surface area (µm) for both using the HALO software (v4.0.5) from immunostaining images of P28 Control and Krt76^-/-^ mice. The mean sum of medullary areas from one section was measured and also used to calculate its proportion to total thymus area, along with the mean number of medulla islets within one thymus.. For each sample, *n*=11-18 sequential sections were analysed from n≥3 biological replicates/group.

For quantification of KRT10^+^ cells, immunostaining images of P28 thymic medulla were analysed using the HALO software (v4.0.5) and HighPlex FL v4.23.214 base algorithm for cell segmentation. Individual cells were segmented using DAPI nuclear staining, and KRT10^+^ cells were selected with a threshold of expression ≤1µm cytoplasmic radius; percentages were calculated compared to total DAPI^+^ cells. To quantify the co-locolisation between KRT10^+^ or Involucrin^+^ cells with KRT76, ROIs on P28 thymus immunostaining images first underwent nuclear segmentation for quantification of total number of cells, followed by quantification of co-localisation or individual expression using ImageJ (v1.54g). Skin and tongue images were quantified using the HALO software (v4.0.5) using the HighPlex FL v4.2.14 algorithm: DAPI nuclear staining was used for cell segmentation and positively stained cells for CD45^+^, CD4^+^, CD8^+^ were quantified; the overall percentage and absolute number of positive cells in total cells (DAPI) was calculated within the skin dermis region or the tongue stromal region. For muscle sections, the absolute number of CD45 positive cells in the field of view was quantified. For quantification of autoantibodies, images were analysed using ImageJ (v1.54g). ROIs encompassing the skin’s epidermis and dermis regions, the tongue’s lamina propria and epithelium, and the squamous stomach epithelium were generated. While for images of muscle sections, full field of view was analysed. The Mean GrayValue was used as a measure of fluorochrome mean intensity. All quantification was performed blinded to genotype or treatment group.

### Preparation of Thymic Populations for scRNA-seq

Thymus was harvested from P28 mice (*n*=4*/*genotype). Individual thymi were crushed in chilled PBS using a pestle to extract the lymphocytes and stored on ice. The remaining stromal tissue was finely sliced and digested enzymatically using 0.05 % [w/v] of Liberase (Roche, 5401119001), 100 U/ml DNase I (Sigma-Aldrich, 10104159001) in RMPI 1640 Medium (ThermoFisher Scientific, 11875093) for 30-40min at 37°C at 100rpm. Cells were mechanically disrupted using a 21G needle to break up aggregates and filtered through at 100µm filter for single cell suspension. Single-cell suspensions were stained for surface markers against: CD45 BV510 (Biolegend, 103138), EpCAM PE (Biolegend, 118206) and DAPI as a live/dead marker (BioLegend 422801) and sorted using FACSAria III cell sorter (FACSDiva 8.0.1 software) for CD45^+^, EPCAM^+^ and CD45^-^EPCAM^-^ as outlined in the gating strategy (**Supplementary Fig. 9b**). The final mixture contained 70% EPCAM^+^ cells, 20% EPCAM^-^CD45^-^ cells, 10% CD45^+^ cells /genotype (Experiment 1). In a separate experiment (Experiment 2), thymi from *n=*8/genotype pooled 4–5-week-old were enzymatically and mechanically dissociated as described above and depleted for CD45-expressing cells via immunomagnetic separation (EasySep Mouse TIL (CD45) Positive Selection Kit, StemCell, 100-0350). The enriched single-cell suspension was stained for: CD45 BV510 (Biolegend, 103138), EpCAM APC (Biolegend, 118213), LY51 PE-Cy7 (Biolegend, 108314), and DAPI as a live/dead marker (BioLegend, 422801) and sorted using FACSAria III cell sorter (FACSDiva 8.0.1 software) for medullary thymic epithelial cells (CD45^-^EPCAM^+^LY51^-^) as described in the gating strategy (**Supplementary Fig. 9b**). In a separate experiment (Experiment 3), *n=*4/genotype pooled P28 thymi were crushed and filtered. The single-cell suspension was stained against CD45 BV510 (Biolegend, 103138), EpCAM PE (Biolegend, 118206), CD4 BUV395 (BDBiosciences, 740208), CD25 APC-Cy7 (eBiosciences, 47-0251-82), CD11c PE (Biolegend, 117348), MHC Class II AF700 (Biolegend, 107622) and DAPI to sort for Tregs (CD4^+^CD25^+^) and DCs (CD45^+^EPCAM^+^CD11c^+^MHC-Class II^+^), the final sample contained an even mixture of the two populations.

### Droplet-based scRNA-Seq, Processing, QC filtering and Integrative Analysis

Where possible 30,000 viable cells were processed to generate libraries for Gene Expression (GEX) Chromium single-cell 5’ reagent v2 protocol as per manufacturer’s instructions. The 10x Genomics libraries were pre-pooled into a single sample and sequenced by NGS Europe (Azenta Life Science Group) or at the Blizard Genome Centre Facility (QMUL, UK) on an Illumina HiSeq 4000, with a sequence configuration of 26-10-10-90 and reads depth of 30,000 read pairs targeted/cell for 10X single-cell from Single Cell 5’ v2 Dual Index for GEX libraries.

The outputs were aligned to the *mus musculus* reference genome (refdata-gex-mm10-2020-A) using the CellRanger 7.0.1 pipeline, according to guidelines outlined by 10x Genomics. The output count files were subsequently filtered and analysed using the *Scanpy* pipeline^57^. Quality control filtering was applied to each sample individually whereby cells exclusion based on <200 genes and <100 counts, genes exclusion based on expression in <3 cells and <10 counts and cells with an *expected_doublet_rate* set between 0.225-0.25 predicted using *Scrublet*^58^. Further thresholds were set roughly to >20% mitochondrial reads, ribosomal reads lower or higher than the 5^th^ and 95^th^ percentile and >1% of hemoglobin reads. Control and Krt76^-/-^ samples were integrated using the *HarmonyPy* algorithm^59^, followed by normalization, log transformation, regression to remove cell-cycle effects and scaling of data.

### Clustering and Sub-Clustering

Dimensionality reduction was calculated using Principal Component Analysis (PCA) (n_pcs=40), followed by UMAP processing using the Leiden algorithm with 13 clusters identified (resolution=0.45) for Experiment 1. The stromal cluster was merged to Experiment 2 which identified 16 clusters (resolution=0.28), cells belonging to one cluster of high counts were removed. Gene cluster markers were used to annotate clusters giving a final 9 clusters. Following annotation, subclustering of all mTECs (mTECsI, Proliferating mTECs, mTECsII, mTECsIII and mTECsIV) or Mimetic mTECs (mTECsIII and mTECsIV) was done to generate new datasets. Processing of the mTECs revealed 14 clusters (resolution=0.38), while the Mimetic mTECs dataset revealed 12 clusters (resolution=0.59). The immune clusters from Experiment 1 were subclustered to reveal 10 clusters (resolution=0.73), the Double-Positive thymocytes and Single-Positive thymocytes barcodes were extracted for V/J analysis. For Experiment 3, processing revealed 14 clusters (resolution=0.6), gene cluster markers were used to annotate which allowed sub-clustering of Tregs-specific clusters for downstream V/J analysis. The GEX dataset DP, SP and Tregs cells from Experiment 1 and Experiment 3 were merged using the function *adata.concat*.

### Identifying *Krt76*-expressing mTECs Signature

The *Krt76* signature was identified in the Mimetic mTECs dataset. In the Control sample only, cells with Krt76 expression >0.05 (using the function *np.where(adata[:,’Krt76’].X>0.05, True, False)*) were compared to those with <0.05 expression via differential gene expression (DEG) analysis using *sc.tl.rank_genes_groups* (method=‘Wilcoxon’). Genes with log fold change≥1.5 were retained. To compare their gene expression between Control and Krt76^-/-^ samples, *Krt76*-expressing cells were extracted using the function: *adata_K76=adata[(adata[:,’Krt76’].X>0.05).flatten(), : ]*.

### mTECs Trajectory & Pseudotime Analysis

We used *Scanpy* and *scVelo* package^60^ (v.0.3.2) for RNA trajectory and pseudotime analysis on the Control and Krt76^-/-^ sample separately from experiment 2 only. Loom files were merged with the annotated .h5ad files. The object underwent quality control filtering and normalization, followed by neighbourhood analysis (n_pcs=40, n_neighbours=30). The splicing kinetics of each gene were computed using *scv.tl.recover_dynamics* from which *scv.tl.velocity* (mode=dynamical) was used to calculate the velocities. mTECsI, Proliferating mTECs, mTECsII and CorneoTECs subsets were extracted for the pseudotime analysis as these represent either less differentiated clusters (mTECsI and Proliferating mTECs) or *Krt76*-expressing clusters (mTECsII and CorneoTECs) and the tool *sc.tl.dpt* was used with the mTECsI cluster set as the root cells based on published reports^14,30^, a list of known markers of epithelial differentiation were selected and plotted across *dpt* pseudotime. For the RNA trajectory analysis, mTECs II and Mimetic mTECs clusters for Control and Krt76^-/-^ sample were extracted and the function *scv.tl.velocity_graph* was used to compute the trajectory across clusters.

### Pseudobulk V/J Feature Comparison

Proportions of V/J gene usage for TCR alpha chains were computed using the scVDJ-seq computational analysis Dandelion^53^. The data was processed using the CellRanger vdj reference (v5.0.0), αβTCR libraries were extracted from the all_contig_annotations.csv outputs. The DP, SP, Tregs merged object outlined above was filtered to only include barcodes with ‘Single pair’, ‘Extra pair’ and ‘Extra pair-exception’ TCR contigs using the default settings of *ddl.tl.vj_usage_pca*, restricting the analysis only to alpha chain V/J genes. The cells were grouped according to combined cell type stage (DPs, SPs and Conv Tregs) in Control and Krt76^-/-^ samples separately prior to performing the calculations. Principal component analysis using the function *sc.pl.pca* was then used to plot the V/J feature space.

### TCRβ Bulk Sequencing and Analysis

Thymus was harvested from 4-6 weeks mice (*n*=4*/*genotype). Individual thymi were crushed in chilled PBS using a pestle to extract the lymphocytes and filtered through a 40µm filter. Single-cell suspensions were stained for surface markers against: CD4 FITC (BD Biosciences, 553729), CD8 BV605 (Biolegend, 100743), CD5 PerCP Cy5.5 (Biolegend, 100624), TCRβ APC (eBioscience, 17-5961-82), CD25 APC Cy7 (eBioscience, 47-0251-82) and DAPI as a live/dead marker (BioLegend, 422801) and sorted using FACSAria III cell sorter (FACSDiva 8.0.1 software) for signalled SP CD4^+^ T cells and non-signalled DP CD4^+^CD8^+^ T cells as outlined in the gating strategy (Supplementary Fig. 9c). TCRβ bulk sequencing was performed through the company ImmunoServ Ltd (https://immunoserv.com/). RNA was extracted and quality checked pior to library preparation using iRepertoire’s long-read dual-index primer (mouse) kits for the TCR β chain (covering the FR2-CDR3 region) following the ‘low input, low quality’ protocol according to manufacturer’s instructions. Pooling was carried out to account for differences in cell numbers between the DP and CD4 samples by pooling at a 2:1 ratio. Sequencing libraries were clustered onto a flowcell on the Illumina Instrument (MiSeq i100, 600 cycles) according to manufacturer’s instructions. The samples were sequenced using a 2×300bp Paired End (PE) configuration. Image analysis and base calling were conducted by the Illumina Control Software. Raw sequence data (.bcl files) generated from the sequencer was converted to fastq files and demultiplexed using Illumina bcl2fasq 2.20 software.

Following sequencing, fastq files were processed using MiXCR (v4.7.0-370-develop) with iRepertoire preset (irepertoire-mouse-rna-xcr-repseq-lr). TCR data from assembled clones in the ‘clones.clns’ files were used for downstream analysis. Only productive TCRs were retained for downstream analyses. For performing machine learning prediction of CD5+ association^61^, TCRs were deduplicated based on V/J call and CDR3 sequence prior to performing prediction. TCR-sequencing data first underwent quality control checking with sc-dandelion (v1.0.0-alpha)^53^ with the ‘check_contigs’ function. We then derived immune repertoire features from each sample using CDR3 amino acid sequences, V and J gene calls, and per-clone UMI counts as a measure of clonal abundance. The distribution of CDR3 lengths per sample was calculated as the UMI-weighted fraction of clones at each length, capturing composition at the level of clone size. CDR3 length was compared directly between experimental groups at the sample level. For each sample, the UMI-abundance-weighted mean CDR3 length was calculated across all clones. Differences in mean CDR3 length across all groups were assessed using the Kruskal-Wallis H test. Pairwise comparisons between groups were subsequently performed using two-sided Mann-Whitney U tests, with resulting p-values corrected for multiple comparisons using the Benjamini-Hochberg FDR procedure. All analyses were performed in Python (v3.13.13) using pandas (v2.3.3) and numpy (v2.4.6) for data handling, scikit-learn (v1.9.0) for distance computation and MDS ordination, scikit-bio (v0.7.3) for PERMANOVA, scipy (v1.17.1) for the Kruskal-Wallis and Mann-Whitney U tests, and statsmodels (v0.14.6) for FDR correction. Figures were generated using matplotlib (v3.10.9).

### Re-analysis of Published Dataset

Results generated using published datasets were used in this study (GEO: E-MTAB-8581, GSE194252, GSE129218, GSE241132)^14,28,31,51,52^. For the analysis of the human and mouse thymus dataset^28^, the .h5ad files downloaded from the E-MTAB-8581 dataset were already filtered, normalised, log transformed and annotated. For both the human and mouse datasets, only the thymic epithelial clusters were extracted and for analysis of the human dataset, only adult samples were used for downstream analysis. For the analysis of the Mimetic mTECs the GSE194252 dataset was used^14^, the 10x Genomics outputs were read using the *sc.read_10x_mtx* to extract cell barcodes and annotation of features. To extract only the adult samples, their annotated metadata file was integrated within the *Scanpy* object. The same QC and filtering steps outlined above were applied. Clustering via the *leiden* algorithm revealed 13 clusters (resolution=0.62); gene cluster markers published in their study were used to annotate clusters. For the analysis of mouse skin gene expression, we used the publicly available dataset GSE129218^31^. Cells in this dataset were pre-filtered by the original authors, but further processing was required, including normalisation, log-transformation, and scaling, as described above. Cluster annotations were already incorporated in the .h5ad files. Dimensionality reduction for visualisation was performed using the *sc.tl.tsne* function. For the analysis of human skin gene expression, we used the publicly available *Skin Explorer*^52^ resource, especially dataset GSE241132^51^.

### Analysis of the expression of TSAs in skin and other organs

The Human Protein Atlas^32^ was used for analysis of protein expression data in human tissues. The website is located at https://www.proteinatlas.org. Gene expression analysis of the different TSAs in the different mouse organs was performed using the public data made available by the Tabula Muris Consortium and the Chan Zuckerberg Biohub^33^.

### Autoantibody profiling

IgG antibody analysis was performed on sera from Control/nu and Krt76^-/-^/nu (*n*=5 animals/genotype combined) using the HuProt Human Proteome Microarray (v4.0, CDI Labs) and processed by PEPperPRINT GmbH (Germany). The microarrays were stained according to the manufacturer’s instructions and the Intensity Difference (a.u.) was calculated by subtracting the signal intensity for each protein to the secondary antibody control signal intensity. The protein list was then filtered by removing any ‘Control’ JHU Clone ID entry, selecting only proteins with greater than or equal to 70% homology UniProt and SwissProt sequence identity to their mouse counterparts in the HuProt v4.0 BLAST Label Curation database and whether they met the Tissue Restricted (TRE) GNF GeneAtlas Specificity as previously described^24^. Following filtering, the intensity difference of these selected proteins was logarithmised for each sample and compared using Log2 Fold Change (Krt76^-/-^/nu / Control/nu). To identify the top proteins of interest, only positive values were retained. Gene Ontology Analysis of Cellular Components (GO_Cellular_Component_2025) was carried out on proteins with a Log2 Fold Change greater than 1.5. Gene expression analysis of the top 31 proteins were plotted in Control and Krt76-/- Mimetic mTECs and in different mouse organs using the public data made available by the Tabula Muris Consortium and the Chan Zuckerberg Biohub^33^.

### Statistical Analysis

All statistical calculations were generated using the GraphPad Prism 10 software. Unless indicated in figure legends, all tests used for calculating statistical significance were either a two-tailed unpaired Student’s *t*-test or multiple *t*-tests (no corrections for multiple comparisons). Significance was determined by *p*-value<0.05 (* *p*<0.05; ** *p*<0.01; *** *p*<0.001; **** *p*<0.0001; *ns p*>0.05). Graphs are either produced in GraphPad Prism 10 or using *Scanpy*, *Scvelo* and *matplotlib* packages for scRNA-seq plots.

## Data and code availability

scRNAseq raw FASTQ, processed counts files and code will be made available to a public repository before the publication of the paper.

